# Evolutionary Diversification of Methanotrophic *Ca*. Methanophagales (ANME-1) and Their Expansive Virome

**DOI:** 10.1101/2022.07.04.498658

**Authors:** Rafael Laso-Pérez, Fabai Wu, Antoine Crémière, Daan R. Speth, John S. Magyar, Mart Krupovic, Victoria J. Orphan

## Abstract

‘*Candidatus* Methanophagales’ (ANME-1) is a major order-level clade of archaea responsible for methane removal in deep-sea sediments through the anaerobic oxidation of methane. Yet the extent of their diversity and factors which drive their dynamics and evolution remain poorly understood. Here, by sampling hydrothermal rocks and sediments, we expand their phylogenetic diversity and characterize a new deep-branching, thermophilic ANME-1 family, ‘*Candidatus* Methanospirareceae’ (ANME-1c). They are phylogenetically closest to the short-chain-alkane oxidizers ‘*Candidatus* Syntrophoarchaeales’ and ‘*Candidatus* Alkanophagales’, and encode ancestral features including a methyl coenzyme M reductase chaperone McrD and a hydrogenase complex. Global phylogeny and near-complete genomes clarified that the debated hydrogen metabolism within ANME-1 is an ancient trait that was vertically inherited but differentially lost during lineage diversification. Our expanded genomic and metagenomic sampling allowed the discovery of viruses constituting 3 new orders and 16 new families that so far are exclusive to ANME-1 hosts. These viruses represent 4 major archaeal virus assemblages, characterized by tailless icosahedral, head-tailed, rod-shaped, and spindle-shaped virions, but display unique structural and replicative signatures. Exemplified by the analyses of thymidylate synthases that unveiled a virus-mediated ancestral process of host gene displacement, this expansive ANME-1 virome carries a large gene repertoire that can influence their hosts across different timescales. Our study thus puts forth an emerging evolutionary continuum between anaerobic methane and short-chain-alkane oxidizers and opens doors for exploring the impacts of viruses on the dynamics and evolution of the anaerobic methane-driven ecosystems.

## Introduction

Anaerobic methanotrophic archaea (ANME) is a polyphyletic group of archaeal lineages that have independently evolved the ability of anaerobic oxidation of methane (AOM), a process that is estimated to globally remove more than 80% of the methane produced in deep-sea sediments^1–4^ by reversing the methanogenesis pathway^5^. Whereas the ANME-2 and ANME-3 lineages share common ancestors with the present-day methanogens of the *Methanosarcinales* order, ANME-1 archaea form their own order *’Candidatus* Methanophagales*’* sister to the non-methane alkane degraders ‘*Candidatus* Syntrophoarchaeales’ and *’Candidatus* Alkanophagales’^6^. Distinctly, ANME-1 can grow beyond the cold and temperate deep-sea habitats that they often share with other ANMEs, uniquely thriving at higher temperatures within hydrothermal environments^5,7–9^. In marine sediments, ANMEs mostly form syntrophic associations with sulfate-reducing bacteria (SRB)^10^ via direct interspecies electron transfer^11,12^. However, some ANME-1 have been observed as single cells or as monospecific consortia without partner bacteria^9,13–18^, and have been proposed to perform hydrogenotrophic methanogenesis^17,18,19^, but physiological experiments have thus far failed to support this hypothesis^12,20^. Overall, it remains largely unclear what factors have contributed to the physiological and ecological diversification of ANME-1 from their short-chain-alkane relatives and other ANME lineages.

Despite the dominance of ANME archaea in many methane-rich ecosystems, viruses targeting ANME lineages are largely unexplored^21–23^. By exploiting and spilling host cellular resources through their replication and lytic cycles, viruses play a major role in the ecological dynamics and nutrient cycling in diverse microbial systems^24^. In deep-sea ecosystems, viral lysis has been estimated to cause annual archaeal mortality that releases up to ~0.3 to 0.5 gigatons of carbon globally^25^. Characterizing the distributions and functions of viruses of ANMEs is thus one of the most important tasks for quantitatively linking ANME physiology to the elemental and energy flows in deep-sea methane-driven ecosystems, as well as understanding the drivers of ANME evolution.

## Results

### Expanded ANME-1 diversity reveals a new deep-branching clade in hydrothermal vents

In this study, we recovered thirteen metagenome-assembled genomes (MAGs) of ANME-1 in native and laboratory-incubated mineral samples from the Southern Pescadero Basin hydrothermal vent system^26^ in the Gulf of California, Mexico (Supplementary Tables 1 and 2). These samples not only expanded the known diversity within the ANME-1a clade, particularly the ANME-1 G60 group^10^, but also contained five novel MAGs and one 1.6Mb circular genome scaffold of a deep-branching clade phylogenetically positioned at the base of the ANME-1 order (Fig.1, Fig. S1, Supplementary Tables 2 and 3). We name this clade ‘*Candidatus* Methanospirareceae’, or ANME-1c, which given its basal position, is the phylogenetically closest ANME-1 to the sister orders of non-methane alkane degraders *Alkanophagales* and *Syntrophoarchaeales*^12^.

**Figure 1.**
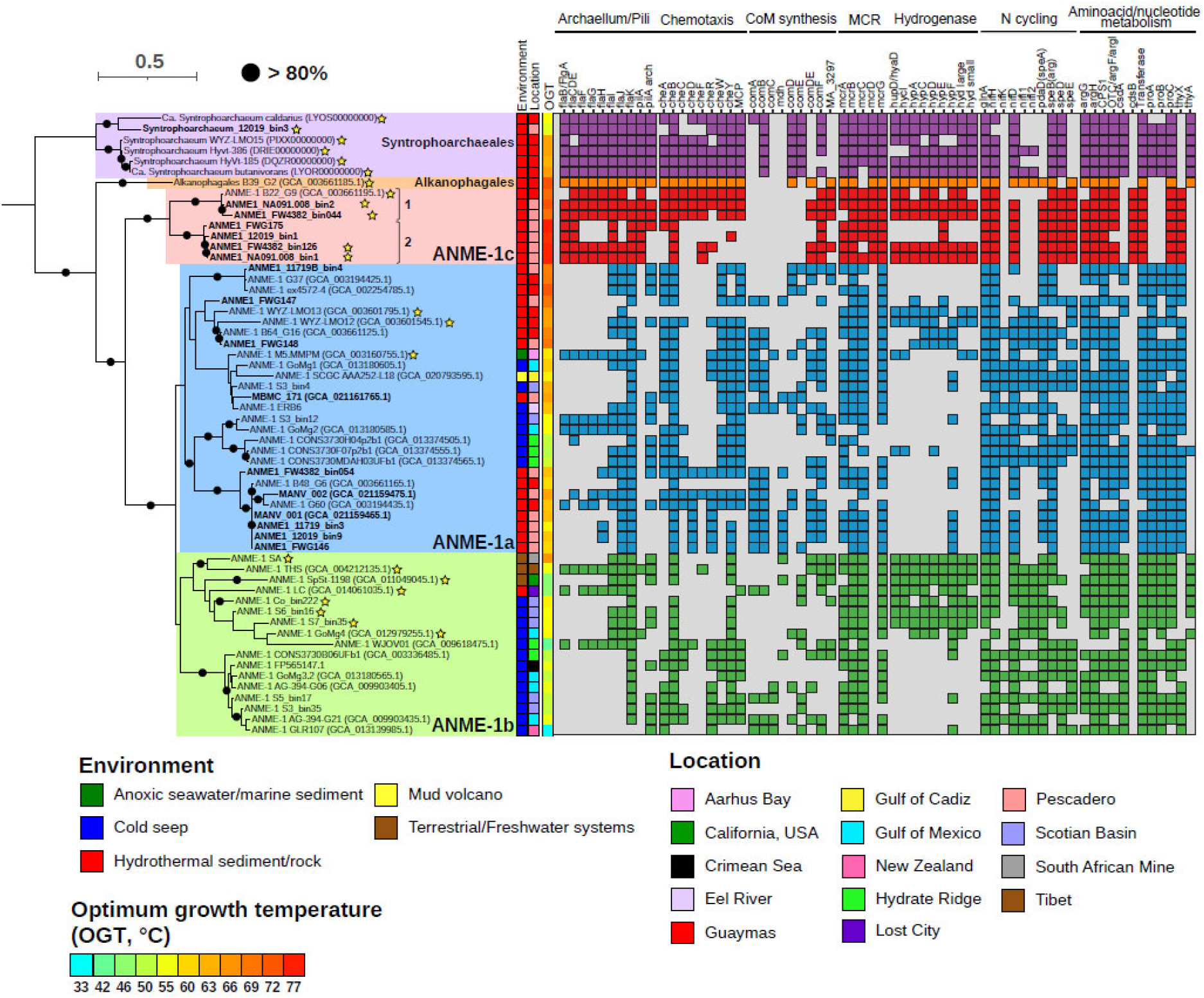
Phylogenomic tree and lineage differentiation of the ANME-1 (*Methanophagales*) order. In bold genomes retrieved from the South Pescadero Basin. The color bars indicate from left to right: environment, location, predicted optimum growth temperature (OGT, in °C) and a genomic comparison of some metabolic features (main text and Supplementary Text). Numbers afther the ANME-1c names indicate the two species of ANME-1c and the stars after the names denote MAGs containing at least the large subunit of a NiFe hydrogenase. Black circles indicate bootstrap support values over 80%. The scale bar represents the number of nucleotide substitutions per site.

Our ANME-1c MAGs represent two different genera ‘*Candidatus* Methanoxibalbensis’ and ‘*Candidatus* Methanospirare’ within the same family with an average nucleotide identity (ANI) of 76%, represented by species ‘*Ca*. Methanoxibalbensis ujae’ (species 1) and ‘*Ca*. Methanospirare jalkutatii’ (species 2, see methods). Based on genome coverage, these two ANME-1 species were the most abundant organisms in rock samples 12019 and NA091.008, while they were hardly detected in rocks 11868 and 11719 and in hydrothermal sediments (Fig.2a).

**Figure 2.**
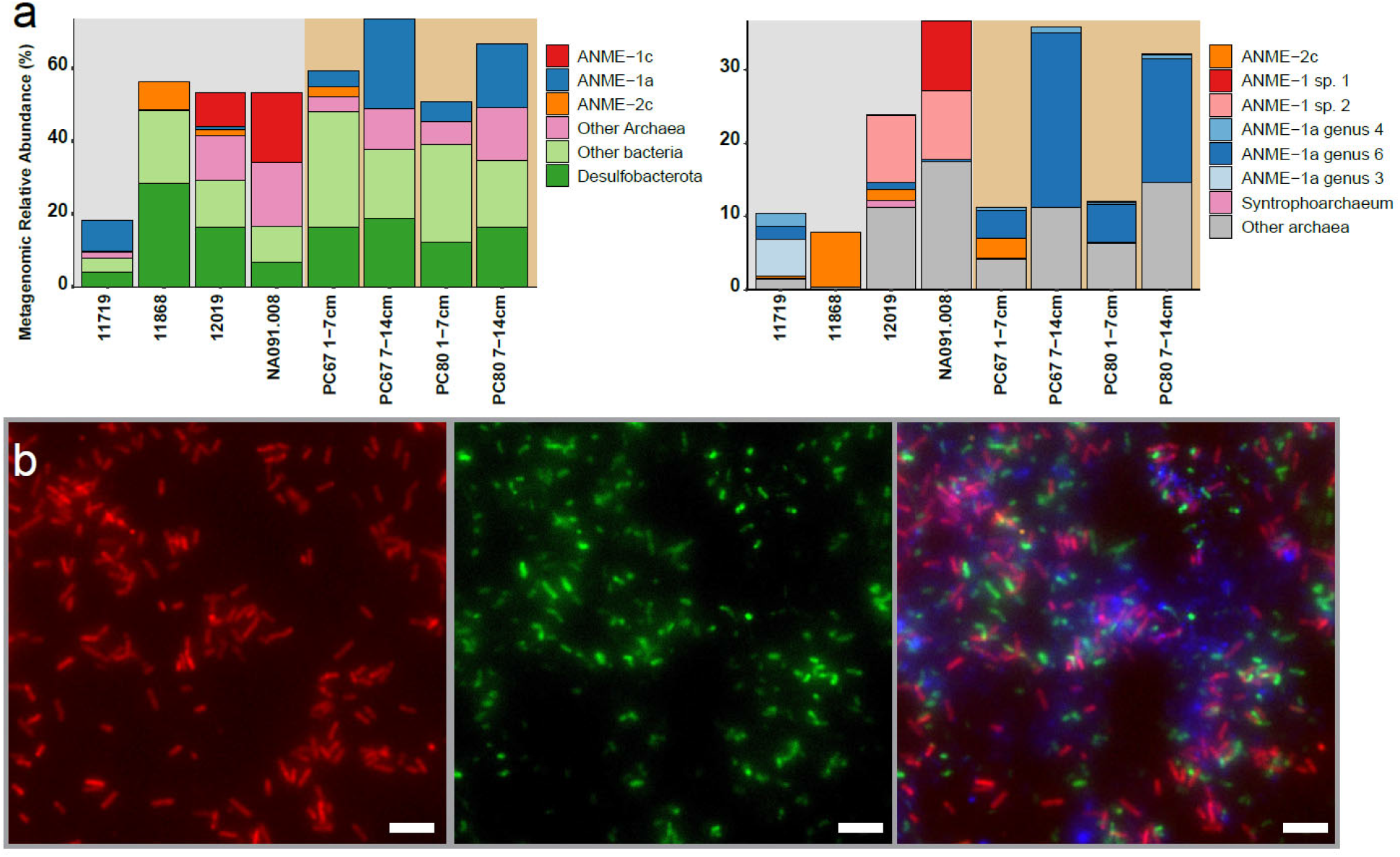
Distribution and morphology of ANME-1 in S. Pescadero sediment and rock samples. **a)** Relative abundance of MAGs from ANME and other bacteria and archaeal lineages (**left**). Genomic abundance for archaea at the species level is on the right, highlighting a variation ANME-1 lineages in rocks and sediments (**right**). Color background indicates rock (grey) or sediment (brown) samples. The total abundance does not reach 100%, since unmapped reads are not included. Note the different scale of the vertical axis between panels. **b)** Fluorescence *in situ* hybridization of ANME-1 cells recovered from rock sample NA091.008. Cells targeted by the general ANME-1-350 probe are shown in red, **left**). Cells targeted by the general bacterial probe 338 are in green, **middle**). A composite overlay showing bacteria (green), ANME-1 cells (red) and DAPI staining of all microbial cells in blue (**right**). Scale bar = 5 μm.

So far, all ANME-1c MAGs and 16S rRNA gene sequences from the NCBI and SILVA databases have originated from hydrothermal environments, specifically the sediments of Guaymas^18,27^ and Southern Pescadero Basins^28^. These hydrothermal habitats are 400 km apart along the same fault system in the Gulf of California and exhibit 20% species-level overlap in the microbial community^28^. This distribution suggests a strong thermophilic physiological specialization of ANME-1c to hydrothermal environments. Indeed, genome-based prediction^29^ suggested a high theoretical optimal growth temperature (OGT, Supplementary Table 4 and Fig.S2) for both ANME-1c species (>70 °C) that was higher than the OGT for both ANME-1a (62 °C) and ANME-1b (52 °C). Such high-temperature adaptation by ANME-1c could be related to their reduced estimated genome size (‘*Ca*. M. ujae’: 1.81 Mb; ‘*Ca*. M. jalkutatii’: 1.62 Mb) as previously observed in other thermophilic bacteria and archaea^30^.

Using fluorescence *in situ* hybridization (FISH) with an ANME-1-targeted 16S rRNA probe, we detected ANME-1 cells in rock NA091.008 (Fig.2b), where ANME-1c were the dominant lineage according to genome coverage (Fig. 2a). These putative ANME-1c cells exhibit the typical cylindrical shape previously reported for other ANME-1 populations^10^ and were loosely associated with bacterial cells in an EPS matrix, or found as single cells.

### Physiological differentiation of diverse ANME-1 archaea

The deep-branching position of ANME-1c led us to examine the genomic patterns of emergence and differentiation of ANME-1 from the sister orders *Alkanophagales* and *Syntrophoarchaeales*. Like all ANME-1, ANME-1c encode a complete reverse methanogenesis pathway including a single operon for the methyl coenzyme M reductase enzyme (MCR), responsible for the activation of methane, and the replacement of F420-dependent methylene-H_4_MPT reductase (Mer) by 5,10-methylenetetrahydrofolate reductase (Met) characteristic for ANME-1^31–33^. Similar to other ANME clades, ANME-1c encodes several multiheme cytochromes (MHC), which likely mediate the transfer of electrons during syntrophic AOM to sulfate-reducing bacteria^5,11,33^.

Notably, ANME-1c exhibit distinct features compared to the ANME-1a and ANME-1b in the operon encoding the MCR enzyme. This enzyme consists of six subunits with the structure α_2_β_2_Y_2_ and the unique nickel-containing cofactor coenzyme F_430_^34^. In the maturation of this cofactor, McrC and McrD, two additional proteins encoded by the MCR operon in methanogens^35^, are involved^36,37^. While *mcrD* is not present in ANME-1a and ANME-1b, both genes are present in ANME-1c, where *mcrD* forms an operon with *mcrABG* (Fig.1). Previous analysis suggested that ANME-1 acquired the *mcr* genes from distant H2-dependent methylotrophic methanogens of the class *Methanofastidiosa*^5^, while they lost the divergent MCRs present in *Syntrophoarchaeales* and *Alkanophagales*, which seem to use larger alkanes. Likewise, we found that the ANME-1c McrD is closely related to the McrD of *Methanofastidiosa* but only distantly related to the McrD of *Syntrophoarchaeales* and *Alkanophagales* that form a different cluster (Fig. S3). These results suggest that during the emergence of ANME-1, a full operon of methane-cycling *mcr* (including *mcrCD)* was acquired by horizontal gene transfer from a *Methanofastidiosa-related* methylotrophic methanogen, and *mcrD* was later lost in both ANME-1a and ANME-1b clades. The ANME-1c also exhibit several additional genomic features that are distinct, highlighted in Fig.1 and described in Supplementary text and Supplementary Table 5.

### Shared origin and differential loss of hydrogenases

Hydrogen was proposed as one of the first candidate intermediates in syntrophic AOM, but fell out of favor after several genomic studies showed that the majority of ANME genomes do not encode hydrogenases. Recent studies however have reported NiFe-hydrogenases in subclades of larger ANME groups, including an ANME-1b subclade ‘*Candidatus* Methanoalium’ and from select ANME-1a genomes (Fig.1)^5,38,39^. Interestingly, the genomes of the sister orders *Syntrophoarchaeales* and *Alkanophagales* encode a NiFe-hydrogenase (Fig. 1)^12^, but physiological experiments discarded a role of this hydrogenase in syntrophic alkane oxidation^12^. Our expanded phylogenomic analysis of ANME-1 confirm that genomes associated with three distinct subclades of the ANME-1a, ANME-1b, and now ANME-1c each encode a NiFe hydrogenase operon (Fig.1). Phylogenetic analysis of the large subunit of these hydrogenases revealed a monophyletic group of ANME-1-affiliated hydrogenases clustering with those of *Syntrophoarchaeales* and *Alkanophagales* (Fig.3a, Supplementary Table 6). Hence, the occurrence of hydrogenases appears to be an ancient trait of the class *Syntrophoarchaeia* that was vertically inherited by the common ancestor of ANME-1 and later differentially lost during ANME-1 clade diversification. Strikingly, the occurrence of hydrogenase has an apparent mosaic distribution among MAGs even within the hydrogenase-containing clades. For instance, within ANME-1c, only two out of five MAGs of *’Ca*. M. jalkutatii’ (FW4382_bin126 and NA091.008_bin1) encode hydrogenases, whereas the complete ’*Ca*. M. jalkutatii’ MAG FWG175, assembled into a single scaffold, does not contain them. To verify that this distribution is caused due to intraspecies variation and not incomplete genome assembly, we conducted independent metagenomic analyses which confirmed the differential presence of hydrogenase genes within ANME strains of different rock samples (Fig. S4, Fig 3b and Supplementary Text). Hydrogenases thus appear to be a part of the pangenomic repertoire of certain ANME-1 subclades/species, likely preserved in the ANME-1 pangenome as an environmental adaptation rather than as an absolute requirement for the methanotrophic core energy metabolism.

**Figure 3.**
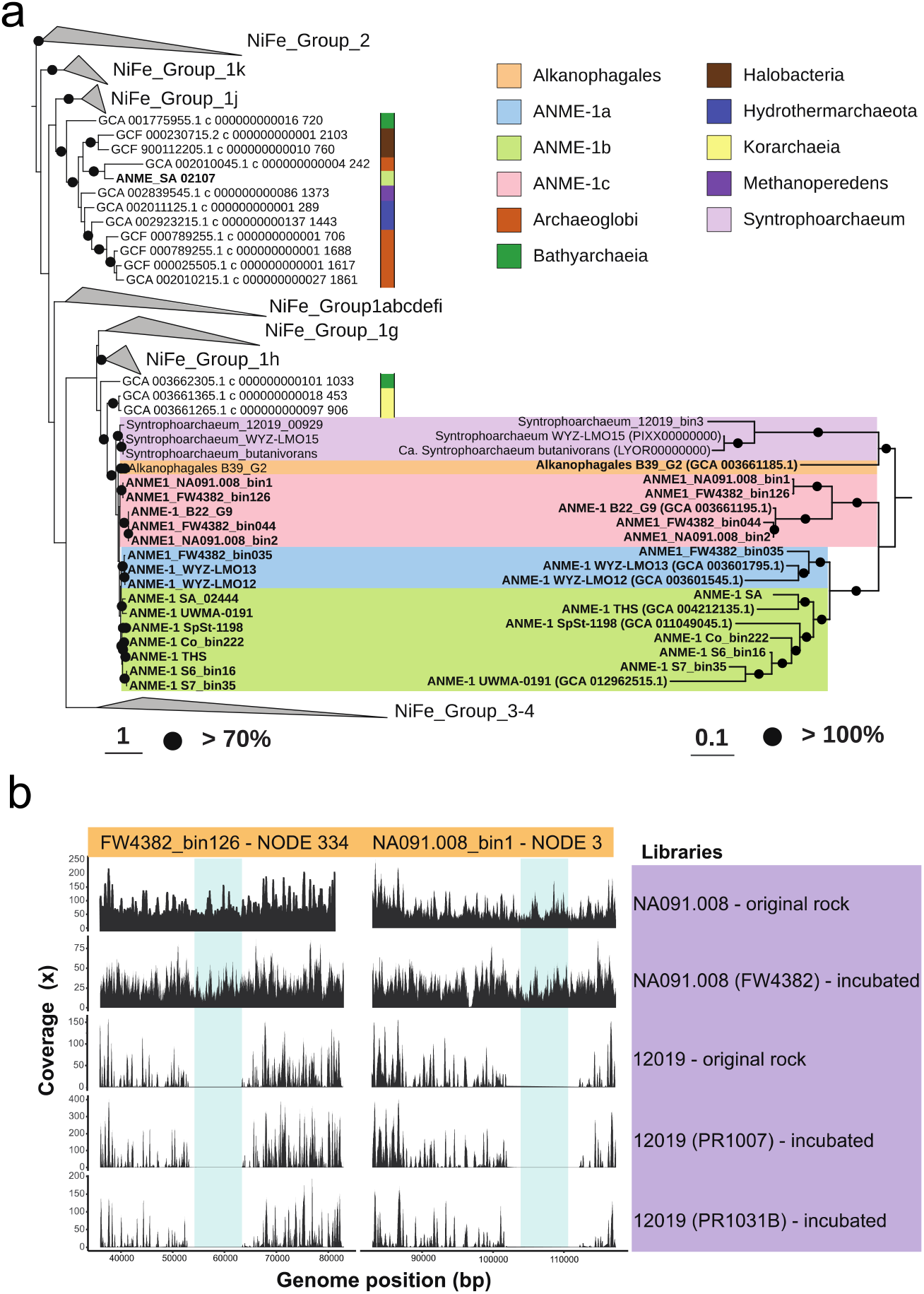
Vertical inheritance and differential loss of hydrogenases across ANME-1. **a)** Phylogenetic tree of the large subunit of the NiFe hydrogenase present in ANME-1 genomes (left) and the corresponding phylogenomic tree of those genomes (right). A few NiFe hydrogenases of ANME-1 genomes were also affiliated with NiFe Group 3 and 4 (not shown, see Supplementary Figure 5). Color bar/backgrounds indicates the phylogenetic affiliation of hydrogenases of interest. Black circles indicate bootstrap support values over 70% (left) and equal to 100% (right). Scale bars represent the number of amino acid substitutions per site. **b)** Read coverage distribution of the hydrogenase operon of ANME-1c genomes FW4382_bin126 and NA091.008_bin1. Metagenomic read libraries are indicated on the right. The blue shade indicates where the hydrogenase operon is located within the corresponding contig.

The potential role of these hydrogenases in ANME-1 is still unclear. Their phylogenetic position, next to hydrogenotrophic enzymes^40,41,42,43^ of the NiFe groups 1g and 1h (Fig. 3a; only a few affiliated to NiFe Group 3 and 4, Fig. S5), suggest a possible involvement in hydrogenotrophic methanogenesis, as previously proposed based on biochemical^44^, environmental^18,19^, isotopic ^17^, and metagenomic data^38^, although enrichment cultivation attempts with hydrogen have been unsuccessful^12,20^. Recently, the genomic analysis of the hydrogenase-encoding ANME-1b group ‘*Ca*. Methanoalium’ showed the presence of distinct electron-cycling features (Rnf complex, cytochrome b) and the absence of multiheme cytochromes suggesting a methanogenic metabolism for this group^5^. By contrast, ANME-1c encodes multiheme cytochromes and lacks these electron-cycling features. Hence, the physiological utility of hydrogenases may vary between lineages. Whereas hydrogen is likely not feasible as the sole intermediate for syntrophic AOM^11,20,45^, it could be produced by ANME-1c as an additional intermediate, as proposed in a mixed model involving direct electron transfer and metabolite exchange^5,46^.

### CRISPR-based discovery of an expansive ANME-1 mobilome

ANME-1 genomes recovered in this study contained various CRISPR-Cas loci (Fig. S6a), enabling the analysis of ANME-1-hosted MGEs through CRISPR spacer-based sequence mapping^47–50^ with additional stringent filters (see Methods and Supplementary Text). Mapping 20649 unique ANME-1 CRISPR spacers to metagenomic assemblies from the Southern Pescadero and Guaymas Basins, and the metagenome-derived virus database IMG/VR v.3^51^ captured 79, 70, and 86 MGE contigs larger than 10 kb, respectively, totaling 235 ANME-1 MGEs (Fig. 4a, S6b). As previously found for the Asgard archaeal mobilome^50^, an apparent cross-site spacer-mobilome mapping indicates a significant fraction of the ANME-1 mobilome has migrated across these sediment-hosted hydrothermal vent ecosystems, along with their hosts^28^ (Fig. 4a). Due to the apparent overlap of CRISPR repeats across diverse ANME-1 lineages, these spacers, and thus the host-MGE interactions, were not further assigned taxonomically to specific ANME-1 subclades. All MGEs identified in this study are distant from all other known viruses (Fig. S7, Supplementary Table 8). A large fraction of these ANME-1 MGEs were found to be interconnected, forming one large complex gene-similarity network of 185 nodes and a medium-sized network of 28 nodes (Fig.4b). The remaining 22 MGEs fell into 7 small groups of 2-3 nodes, and 7 singletons.

**Figure 4.**
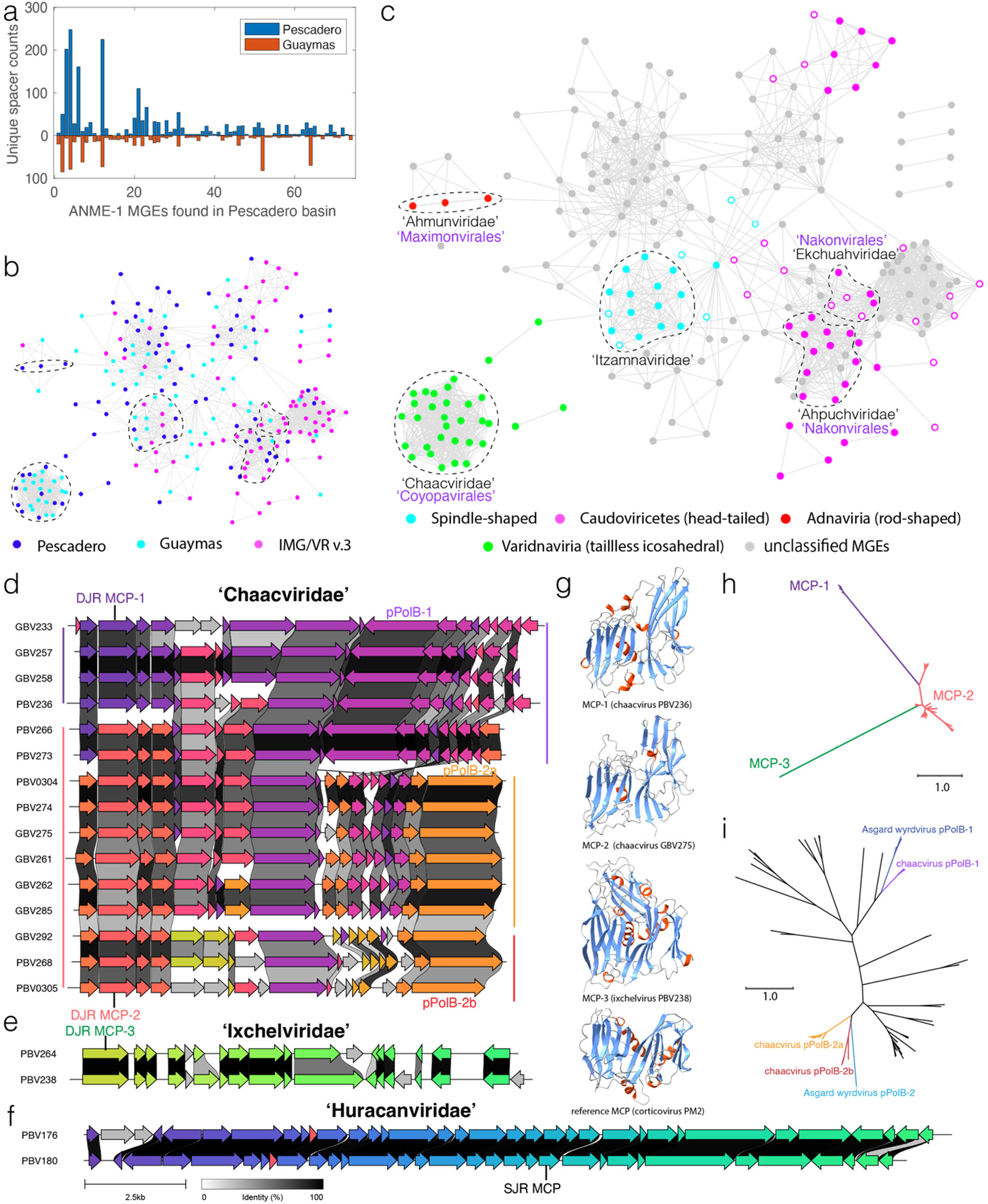
Expansive ANME-1 mobilome includes 16 new viral families and new types of major capsid proteins. **a**) Histograms showing the number of CRISPR spacers from S. Pescadero and Guaymas basin metagenomes matching the S. Pescadero ANME-1 MGEs. **b)**. Gene sharing network of diverse ANME-1 MGEs of different origins. **c**, ANME-1 MGEs, exhibited in the same network as b, are found to encompass major archaeal virus diversity and non-viral elements. Solid/open circles indicate viral assemblies with/without identifiable MCPs. In **b** and **c**, dashed lines encircle five new viral families containing complete genome representatives. The newly proposed names of viral families (black) and orders (purple) are indicated in c. **d-f**. Gene synteny of three new families of tailless icosahedral viruses targeting ANME-1. Different colors indicate 83 different protein groups. The scale bar and precent identity shading are indicated in f. **g**. alphafold2-predicted structures of newly discovered DJR MCPs in ANME-1 viruses shown in panels d and e. Blue indicates β barrels, and red α helices. **h.** Maximum-likelihood analysis of new MCP families indicates their long evolutionary distances. **i**. Maximum-likelihood analysis of PolB found in different clades of the *Chaacviridae* are related to two clades of spindle-shaped Wyrdviruses targeting Asgard archaea. Abbreviations: MCP, major capsid protein. DJR, double jellyroll. SJR, single jellyroll. pPolB, Protein-primed DNA Polymerase family B.

Based on the conservation of signature genes encoding viral structural proteins, we concluded that these MGEs encompass double-stranded DNA viruses belonging to at least 4 widely different virus assemblages characterized by different evolutionary histories and distinct virion morphologies (Fig. 4c). In particular, head-tailed viruses of the class *Caudoviricetes* (realm *Duplodnaviria)* encode characteristic HK97-fold major capsid proteins (MCP) as well as the large subunit of the terminase and portal proteins^52,53^; tailless icosahedral viruses of the realm *Varidnaviria* are characterized by double jelly-roll MCPs ^52,54,55^; viruses of the realm *Adnaviria* encode unique α-helical MCPs which form claw-like dimers that wrap around the viral DNA forming a helical, rod-shaped capsid^56–58^; and all spindle-shaped viruses encode unique, highly hydrophobic α-helical MCPs^59,60^ (Supplementary Table 9-10). In total, 16 new candidate viral families were discovered in this study, including five families with representative complete genomes (Fig. 4c). We named these candidate virus families after Mayan gods, owing to their discovery in the Gulf of California hydrothermal vents off the coast of Mexico (see Methods for etymology of virus family names).

### Tailless icosahedral ANME-1 viruses with previously undescribed major capsid proteins

Tailless icosahedral viruses (*Varidnaviria*) infecting ANME-1 are well distinguished from known viruses, with all 32 representatives unique to this study. They form three disconnected modules, and based on gene similarity analysis, represent three new viral families (Fig. 4c–f)). Members of the ‘Huracanviridae’ encode single jelly-roll (SJR) MCPs related to those conserved in the kingdom *Helvetiavirae*, whereas ‘Chaacviridae’ and ‘Ixchelviridae’ do not encode MCPs with sequence homology to other known viruses. However, structural modeling of the candidate MCPs conserved in ‘Chaacviridae’ and ‘Ixchelviridae’ using AlphaFold2^61^ and RoseTTAFold^6^ revealed unambiguous similarity to the MCPs with a double jelly-roll (DJR) fold (Fig.4g). Phylogenetic analysis revealed that these DJR MCPs form three highly divergent groups, MCP-1-3 (Fig.4g, h), with MCP-2 and MCP-3 containing an additional small beta-barrel which is predicted to point outwards from the capsid surface and likely mediate virus-host interactions.

‘Chaacviridae’ have linear dsDNA genomes with inverted terminal repeats (ITR) and, accordingly, encode protein-primed family B DNA polymerases (pPolB). Chaaviruses display a remarkable genome plasticity – not only do these viruses encode two different variants of the DJR MCPs, MCP-1 and MCP-2, but also their pPolBs belong to two widely distinct clades. Notably, the two MCP and two pPolB variants do not strictly coincide, suggesting multiple cases of recombination and gene replacement within the replicative and morphogenetic modules (Fig. 4d). Maximum likelihood analysis of these divergent groups of pPolB sequences revealed relatedness to two separate clades of pPolBs encoded by Wyrdviruses, spindle-shaped viruses that target Asgard archaea^62^ (Fig. 4i). In addition to pPolB, upstream of the MCP gene, all chaacviruses encode a functionally uncharacterized protein with homologs in Asgard archaeal viruses of the Huginnvirus group, where they are also encoded upstream of the MCP genes^62,63^. This observation suggests a remarkable evolutionary entanglement between these ANME-1 and Asgard archaeal viruses, potentially facilitated by the ecological (i.e., deep-sea ecosystems) rather than evolutionary proximity of the respective hosts.

### Complex ANME-1 viruses with unique structural and replicative features

The head-tailed viruses targeting ANME-1 encode the typical morphogenetic toolkit shared between all members of the *Caudoviricetes*, including the HK97-fold MCP, portal protein, large subunit of the terminase and various tail proteins^58,64^. MCP phylogeny indicates a shared ancestry for the structural components of the viruses of ANME-1 and haloarchaea, which are related at the phylum level (Fig. S8). However, global proteome-based phylogeny^65^ revealed a clear division between ANME-1 and haloarchaeal head-tailed viruses (Fig. 5a). This result suggests that although these viruses encode related core proteins for virion formation, as suggested by their interspersed MCP phylogenetic positions (Fig. S8), the overall proteome contents of ANME-1 and haloarchaeal viruses differ considerably, likely reflecting the adaptation to their respective hosts and ecological contexts. Based on the minimum genetic distances between halovirus families and cross-genome comparisons (Fig. S9), we propose nine new candidate *Caudoviricetes* families. Viruses in these new families exhibit little proteome overlap with each other (Fig. S9), further illustrating the vast genetic diversity of ANME-1 head-tailed viruses.

**Figure 5.**
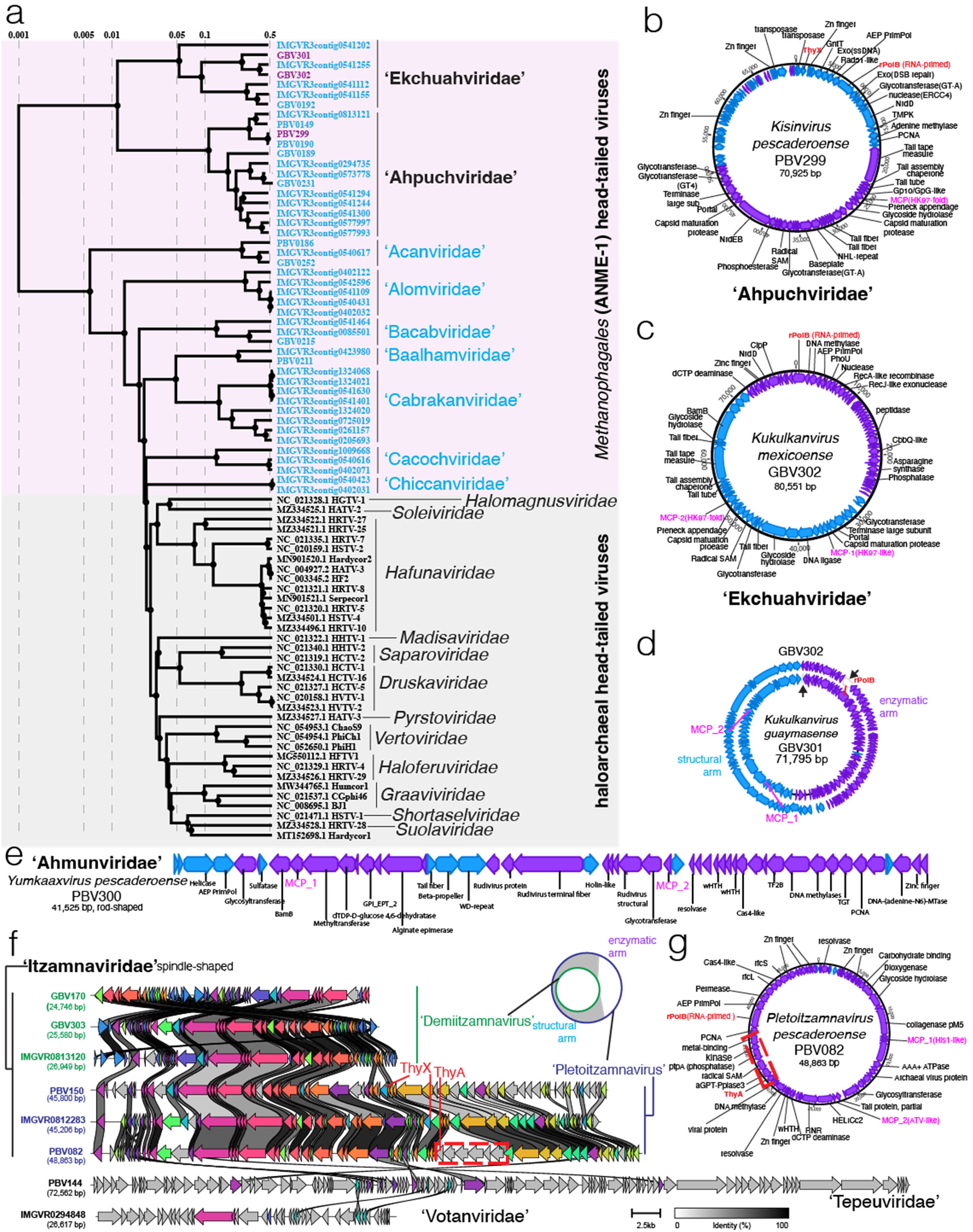
ANME-1 viral genomes encode complex structures. **a**) Evolutionary division between headtailed viruses targeting ANME-1 and haloarchaea revealed by global proteome-based phylogenetic analyses. ANME-1 viruses with complete circular genomes are highlighted in purple, those with unconfirmed completeness are in blue. **b)** and **c)** genome organization and gene content of the complete genomes representing two new families of ANME-1 head-tailed viruses. Blue and purple shading represents forward and reverse strands, respectively. MCP, PolB and ThyX genes are highlighted in pink and red. **d**) circular alignment of the two genomes of *Ekchuahviruses*. Black arrowheads indicate the original contig start/end sites in each assembly. **e**) Gene content of the complete linear genome of a representative of the rod-shaped virus family *Ahmunviridae*. **f**) Gene synteny of three new families of spindle-shaped viruses targeting ANME-1, where complete, circularized genomes of Itzamnaviridae were found to occur in two genome sizes, where Demiitzamnavirus representatives align with a section of the larger Pletoitzamnavirus genomes (illustrated on the top-right). Different colors indicate 76 different protein groups. Grey shading denotes singletons. The scale bar and percent identity shading are indicated in the bottom right. **g**) Gene content of the complete linear genome of a representative of the spindleshaped virus family *Itzamnaviridae*. Dashed red box in f and g highlights an example of a multi-gene cluster insertion. In d and f, structural arm denotes the genome fraction where all viral structural genes reside; enzymatic arm denotes the fraction where no structural genes but only enzyme-encoding genes reside.

‘Ekchuahviridae’ and ‘Ahpuchviridae’ are represented by ANME-1 viruses with complete 70-80 kb genomes and in the proteomic tree form sister clades outside of the 3 orders of haloviruses, forming a new order ‘Nakonvirales’ (Fig. 5a). The Southern Pescadero Basin ahpuchviruses PBV299 (70.9 kb, complete, Fig. 5b) and IMGVR0573778 (74.8 kb, near complete) each encode one copy of MCP, while the two Guaymas Basin ekchuahviruses, GBV302 (80.6 kb, complete, Fig. 5c) and GBV301 (71.8 kb, complete)^51,66^, each encode two MCP copies. This is unique among other known *Caudoviricetes* targeting haloarchaea and ANME-1. We can exclude an assembly artifact, as the initial assemblies of the two ekchuahviruse*s* were found to have a circular alignment with each other (Fig. 5d). Both MCP genes are accompanied by cognate capsid maturation protease genes, whereas all other virion morphogenetic proteins are encoded as single copy genes (Fig. 5c). MCP-1 is likely ancestrally conserved, whereas MCP-2 appears horizontally transferred from haloferuviruses. Their large phylogenetic distance suggests a long co-existence and co-evolution in ekchuahviruses.

The coexistence of two divergent MCP genes is also found in members of putative rod-shaped viruses within the new family ‘Ahmunviridae’, which we propose including into the class *Tokiviricetes* (realm *Adnaviria*) within a new monotypic order ‘Maximonvirales’ (Fig. 5e), and viruses with predicted spindle-shaped morphology, the ‘Itzamnaviridae’ (Fig. 5f, g). These two new clades of viruses are represented by complete linear genomes with inverted terminal repeats and circular genomes, respectively. This contrasts another spindle-shaped ANME-1 virus, the tepeuvirus PBV144, which has the largest genome (72.6 kb, not yet circularized) but only one MCP. The coexistence of divergent MCPs is unusual among *Caudoviricetes*, but has been previously documented for the head-tailed T4 phage, whose MCPs respectively form hexameric and pentameric capsomers occupying the 5-fold icosahedral vertices^67^. Dual-MCP rod-shaped viruses either form a functional MCP heterodimer^56,57,68^ or use only one copy for virion formation^57,58^. It is thus yet unclear how coexisting MCP genes contribute to the capsid architecture of ANME-1 viruses.

### Pan-virus auxiliary functions and virus-driven ANME-1 evolution

The large genomes of head-tailed and spindle-shaped viruses of ANME-1 exhibit strong clustering of functionally related genes: one half of the viral genome contains all structural genes, while the other half encodes diverse enzymes involved in DNA synthesis and modification and various metabolic and defense functions (Fig. 5b-d,f,g). Notably, the entire ~20-kb replicative/metabolism module is missing from the circular genomes of demiitzamnaviruses. Cross-genome alignments revealed a larger variation in gene content for the enzymatic arms in both head-tailed and spindle-shaped viruses, frequently in the form of multi-gene cluster insertions (Fig. 5f, Fig. S9). Head-tailed ‘Ekchuahviridae’ and ‘Ahpuchviridae’ and spindle-shaped ‘Pletoitzamnavirus’ and ‘Tepeuviridae’ encode RNA-primed family B DNA polymerases, which are commonly encoded by dsDNA viruses with larger genomes^69^. The structural-enzymatic arm split thus resembles the core- and pan-genomes of microbes, allowing versatile interactions between these viruses and their ANME-1 hosts (Supplementary Table 10). For example, head-tailed and spindle-shaped viruses contain auxiliary metabolic genes (AMGs) involved in nucleotide and amino acid metabolisms (NrdD, QueCDEF, and asparagine synthase), carbon anabolism (PEPCK and GntT), and phosphorous and sulfur anabolism (PhoU and PAPS) (See Supplementary Table 11 and Supplementary text for details).

Our analysis of viral AMGs also suggested the involvement of viruses in the ancestral metabolic diversification of ANME-1. Specifically, the detection of genes encoding ThyX, which catalyzes dUMP methylation into dTMP and likely boosts host thymidine synthesis during viral production, in head-tailed ahpuchviruses and ekchuahviruses and in spindle-shaped pletoitzamnaviruses (Fig. 5b, 5f). This coincides with the presence of *thyX* in the ANME-1 host, which unlike other ANME lineages and short-chain-alkane-oxidizing archaea, do not encode the non-homologous thymidylate synthase gene, *thyA*^5^ (Fig 1). The dichotomous distribution of the functional analogs *thyA/thyX* is prevalent across microbes and notably, in itzamnaviruses *thyX* and *thyA* may exist in different members (Fig. 5f, 5g). Phylogenetic analysis of the ThyX show that the ThyX encoded by ANME-1 and their viruses form a distinct clade distant from those encoded by bacteria, archaea and other *Caudoviricetes* (Fig. 6, Fig. S10a, S10b). Strikingly, ThyX encoded by itzamnaviruses form a well-supported monophyletic group located at the base of this divergent clade, and the deep-branching ANME-1c encode ThyX that belong to the second deepest branch. Notably, the Guaymas Basin ANME-1c bin B22_G9 contains both a genomic *thyX*, as well as *thyX* encoded by a partial itzamnavirus-derived provirus. Ekchuahviruses and ahpuchviruses likely acquired *thyX* independently at a later stage (Fig. 6, Fig. S10a, S10c).

**Figure 6.**
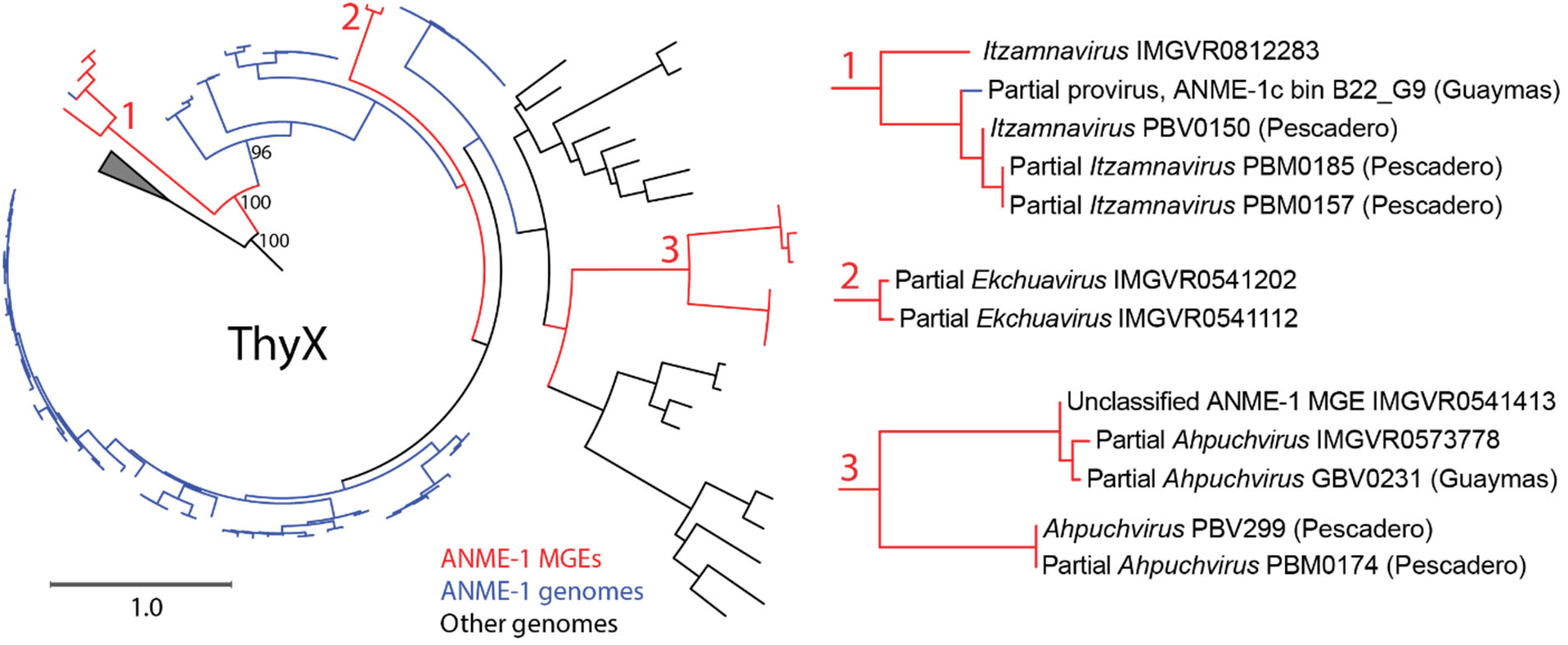
A viral origin of thymidylate synthase in ANME-1. Maximum likelihood analysis of ThyX related to ANME-1 encoded ThyX proteins, with expanded views of ThyX from ANME-1 viruses on the right. Legends for the branch colors for ThyX from mobile genetic elements (MGEs) and ANME-1 genomes are indicated below the main phylogenetic tree.

The above analyses suggest that *thyX* was first acquired by spindle-shaped ANME-1 viruses and then transmitted into the common ancestors of ANME-1, displacing *thyA*. Due to higher promiscuity of viral DNA polymerases and the intense arms race, viral genes are known to evolve rapidly^70^, which is in line with the extreme divergence of the ANME-1/viral *thyX* from the canonical clade.

## Discussion

In this study, metagenomic characterization of a new hydrothermal vent environment in the Southern Pescadero Basin led to the expansion of the known ‘*Ca*. Methanophagales’ (ANME-1) diversity to include ‘*Ca*. Methanospirareceae’ (ANME-1c) and their viruses. ‘*Ca*. Methanospirareceae’ is a deep-branching family that so far has only been detected in high temperature hydrothermal environments. Comparative genomics indicates an evolutionary continuum within the class *Syntrophoarchaeia*, as ANME-1c retained various ancestral features also found in ‘*Ca*. Syntrophoarchaeales’ and *’Ca*. Alkanophagales’, including hydrogenases. The phylogeny of these hydrogenases is congruent with the genome phylogeny indicating an apparent vertical inheritance and differential loss of these genes in ANME-1, suggesting that these hydrogenases have a nonobligatory physiological role, but may confer a long-standing selective advantage.

Our study also uncovered a putative viral source of the ANME-1-specific thymidylate synthase gene *thyX*that replaced the functional analog *thyA* gene. ThyX differs from ThyA in its use of NADPH as the electron donor when transferring the methyl group from the C1 intermediate H_4_MPT=CH_2_ to dUMP to yield dTMP, without oxidizing the H4MPT moiety^5^. H_4_MPT is a core co-factor constantly recycled through the Wood-Ljungdahl pathway that fuels ANME-1 anabolism^5,71^; NADPH abundance is highly dependent on the type of host energy metabolism and redox state^72,73^. The virus-induced ThyA-to-ThyX transition may have played a role in the metabolic diversification and subsequent ecological expansion of the ANME-1 ancestors. C_1_ anabolism appears to be more divergent across ANME lineages than their C_1_ energy metabolism^5^, which may have also originated from viruses and other MGEs as well.

The expansive virome of ANME-1, as discovered in this study, is distant from all known viruses, forming 16 previously undescribed families and at least 3 new orders. They are characterized by many unique structural and replicative features, significantly expanding our appreciation of the archaeal virus diversity and their ecological significance. These findings open doors for targeted culture-dependent and culture-independent exploration of ANME virus-host interactions that are expected to play a critical role in the biogeochemical cycling^25,74^ in these productive methane-driven ecosystems^1–4^.

## Material and Methods

### Sampling and incubations

Four samples of minerals were collected from the 3.7 km-deep Auka vent field in the Southern Pescadero Basin (23.956094 N, 108.86192 W)^26,28,75^. Sample NA091.008 was collected and incubated as described previously^50^. Samples 12019 (S0200-R1), 11719 (S0193-R2) and 11868 (S0197-PC1), the latter representing a lithified nodule recovered from a sediment push core, were collected with *ROV SuBastian* and *R/V Falkor* on cruise FK181031 in November 2018. These samples were processed shipboard and stored under anoxic conditions at 4 °C for subsequent incubation in the laboratory. In the laboratory, mineral sample 12019 and 11719 were broken into smaller pieces under sterile conditions, immersed in N_2_-sparged artificial sea water and incubated under anoxic conditions with methane as described previously for NA091.008^50^. Additional sampling information can be found in Supplementary Table 1. Mineralogical analysis by XRD identified several of these samples as containing barite (11719, NA091.008), collected from two locations on the western side of the Matterhorn vent, and one sample (12019) recovered from the sedimented flanks from the southern side of Z vent, which was saturated with oil. Our analysis also includes metagenomic data from two sediment cores (DR750-PC67 and DR750-PC80) collected in April 2015 with the *ROV Doc Ricketts* and *R/V Western Flyer* (MBARI2015), previously published^28^.

### Fluorescence *in situ* hybridization

Samples were fixed shipboard using freshly prepared paraformaldehyde (2 vol% in 3x PBS, EMS) at 4°C overnight, rinsed twice using 3x PBS, and stored in ethanol (50% in 1xPBS) at −20°C until processing. Small pieces (< 1cm^3^) of the mineral sample NA091.008 were gently crushed in a sterile agate mortar and pestle in a freshly prepared, filter sterilized 80% ethanol - 1× PBS solution. About 500 μl of the resulting mixture was sonicated three times in 15 second bursts on a Branson Sonifier W-150 ultrasonic cell disruptor (level 3) on ice with a sterile remote-tapered microtip probe inserted into the liquid. Cells were separated from mineral matrix using an adapted protocol of Percoll density separation^11^. The density-separated cells were filtered onto 25 mm polycarbonate filters with a pore size of 0.22 μm, and rinsed using 1x PBS. Fluorescence *in situ* hybridizations were carried out as described previously^11^ using a 1:1 mixture of an ANME-1 targeted probe (ANME-1-350^14^ labeled with Cy3) and the general bacterial probe mix (EUB-338 I-III^76,77^ labeled with Alexa-488) at 35% of formamide concentration. Hybridized samples were imaged using a 100x objective using a Zeiss Elyra structured illumination microscope using the Zen Black software.

### DNA extraction and sequencing

DNA extraction from the mineral samples followed previously published protocols^50^. Metagenomic analysis from the extracted genomic DNA was outsourced to Quick Biology (Pasadena, CA, USA) for library preparation and sequencing. Libraries were prepared with the KAPA Hyper plus kit using 10 ng of DNA as input. This input was subjected to enzymatic fragmentation at 37°C for 10 min. After end repair and A-tailing, the DNA was ligated with an IDT adapter (Integrated DNA Technologies Inc., Coralville, Iowa, USA). Ligated DNA was amplified with KAPA HiFi HotStart ReadyMix (2x) for 11 cycles. Post-amplification cleanup was performed with 1x KAPA pure beads. The final library quality and quantity were analyzed and measured by Agilent Bioanalyzer 2100 (Agilent Technologies, Santa Clara, CA, USA) and Life Technologies Qubit 3.0 Fluorometer (Life Technologies, Carlsbad, CA, USA) respectively. Finally, the libraries were sequenced using 150 bp paired-end reads on Illumina HiSeq4000 Sequencer (Illumina Inc., San Diego, CA). After sequencing, primers and adapters were removed from all libraries using bbduk^78^ with mink=6 and hdist=1 as trimming parameters, and establishing a minimum quality value of 20 and a minimal length of 50 bp. For incubated samples, DNA was amplified using multiple displacement amplification (MDA) with the QIAGEN REPLI-g Midi kit prior to library preparation for nanopore sequencing. Oxford Nanopore sequencing libraries were constructed using the PCR-free barcoding kit and were sequenced on PromethION platform by Novogene Inc.

### Metagenomic analysis

The sequencing reads from unincubated rocks were assembled individually and in a coassembly using SPAdes v. 3.12.0^79^. From the de-novo assemblies, we performed manual binning using Anvio v. 6^80^. We assessed the quality and taxonomy affiliation from the obtained bins using GTDB-tk^81^ and checkM ^82^. Genomes affiliated to ANME-1 and *Syntrophoarchaeales* were further refined via a targeted-reassembly pipeline. In this pipeline, the original reads were mapped to the bin of interest using bbmap, then the mapped reads were assembled using SPAdes and finally the resulting assembly was filtered discarding contigs below 1500 bp. This procedure was repeated during several rounds (between 11-50) for each bin, until we could not see an improvement in the bin quality. Bin quality was assessed using the checkM and considering the completeness, contamination (< 5%), N50 value and number of scaffolds. The resulting bins were considered as metagenome-assembled genomes (MAGs). The sequencing reads for the incubated rocks 12019 and 11719 were assembled as described previously for NA091.R008^50^. Additionally, the assembly of 12019 was then scaffolded using Nanopore reads through two iterations of LRScaf v1.1.10^83^. The final assemblies were binned using metabat2 v2.15^84^ using default setting. Automatic metabolic prediction of the MAGs was performed using prokka v. 1.14.6^85^ and curated with the identification of Pfam^86^ and TIGRFAM^87^ profiles using HMMER v. 3.3 (hmmer.org); KEGG orthologs^88^ with Kofamscan^89^ and of COGs^90^ and arCOGs motifs^91^ using COGsoft^92^. To identify multiheme cytochromes in our genomes, we searched the motif CXXCH across the amino acid sequences predicted for each MAG. Similar metabolic predictions were carried out with publicly available ANME-1 and *Syntrophoarchaeales* genomes in order to compare the metabolic potential of the whole ANME-1 order. A list of the genomes used in this study can be found in Supplementary Table 2. For the comparison of different genomic features among the ANME-1 genomes, we searched for specific proteins using the assigned COGs, arCOGs and KEGG identifiers (Supplementary Table 5).

### Genomic relative abundance analysis

We used the software coverM v. 0.5 (https://github.com/wwood/CoverM) to calculate the genomic relative abundance of the different organisms of our samples using all the MAGs we have extracted from our metagenomic analysis. We ran the software with the following additional parameters for dereplication (“--dereplication-ani 95 -- dereplication-prethreshold-ani 90-dereplication-precluster-method finch”). Results were visualized in R^93^ using ggplot ^94^.

### Optimum growth temperature analysis

We calculated the optimum growth temperature for all ANME-1 and *Syntrophoarchaeales* MAGs included in our analysis (Supplementary Table 2) using the OGT_prediction tool described in Sauer and Wang (2019)^29^ with the regression models for Archaea excluding rRNA features and genome size.

### Analysis of the hydrogenase operon of ‘*Candidatus* Methanospirare jalkutatii’ genomes

Since only two of the five genomes of ‘*Ca*. Methanospirare jalkutatii’ have an operon encoding a hydrogenase, we performed additional analysis to better understand this intraspecies distribution. On the one hand, we mapped the metagenomic reads from samples with genomes of ‘*Candidatus* Methanospirare jalkutatii’ (12019, FW4382_bin126, NA091.008, PR1007, PR1031B) to the MAGs containing the hydrogenase operon (FW4382_bin126, NA091.008_bin1) to check if reads mapping this operon are also present in samples from where MAGs without the hydrogenase were recovered. For mapping the reads, we used bowtie2^95^ and then transformed the sam files to bam using samtools^96^ and finally extract the coverage depth for each position. Additionally, we performed a genomic comparison of the genomes with a hydrogenase operon (FW4382_bin126, NA091.008_bin1) with the genome FWG175 that was assembled into a single scaffold. For this, we used the genome-to-genome aligner Sibelia^97^ and we visualized the results using Circos^98^.

### Phylogenetic analysis

For the phylogenomic tree of the ANME-1 MAGs, we used the list of genomes present in Supplementary Table 2. As marker genes, we used 31 single copy genes (Supplementary Table 5) that we extracted and aligned from the corresponding genomes using anvi-get-sequences-for-hmm-hits from Anvio v. 6 ^80,99^ with the parameters “--return-best-hit--max-num-genes-missing-from-bin 7 -- partition-file”. Seven genomes missed more than 7 marker genes and were not used for the phylogenomic reconstruction present in Figure 1 (ANME-1 UWMA-0191, Syntrophoarchaeum GoM_oil, ANME-1 ERB7, ANME-1 Co_bin174, ANME-1 Agg-C03, PB_MBMC_218, FW4382_bin035). The concatenated aligned marker gene set was then used to calculate a phylogenomic tree with RAxML v. 8.2.12^100^ using a partition file to calculate differential models for each gene the following parameters “-m PROTGAMMAAUTO-f a - N autoMRE-k”. The tree was then visualized using iTol^101^. For the clustering of the MAGs into different species, we dereplicated the ANME-1 MAGs using dRep v. 2.6.2 with the parameter “-S_ani 0.95”^102^. A smaller phylogenomic tree was calculated with the genomes containing hydrogenase genes (Fig. 3). For this tree we also used Anvio v. 6 and RAxML v. 8.2.12 with the same parameters but excluding the flag “— max-num-genes-missing-from-bin” from the anvi-get-sequences-for-hmm-hits command to include in the analysis those genomes with a lower number of marker genes that still contain hydrogenase genes (PB_MBMC_218, FW4382_bin035, ANME-1 UWMA-0191).

The 16S rRNA gene phylogenetic tree was calculated for the 16S rRNA genes predicted from our genome dataset that were full-length. We included these full-length 16S rRNA genes in the SILVA_132_SSURef_NR99 database^103^ and with the ARB software^104^ we calculated a 16S phylogenetic tree using the maximum-likelihood algorithm RAxML with GTRGAMMA as the model and a 50% similarity filter. One thousand bootstrap analyses were performed to calculate branch support values. The tree with the best likelihood score was selected.

For the construction of the hydrogenase phylogenetic tree ((Supplementary Table 6), we used the predicted protein sequence for the large subunit of the NiFe hydrogenase present in the genomes of our dataset (Supplementary Table 2), a subset of the large subunit hydrogenases present in the HydDB database^43^ and the predicted hydrogenases present in an archaeal database using the COG motif for the large NiFe hydrogenase (COG0374) with the Anvio v. 6 software. For the mcrD gene phylogeny, we used the predicted protein sequences of mcrD in the ANME-1c genomes and in the previously mentioned archaeal database with the TIGR motif TIGR03260.1 using also the Anvio v. 6 software. The list of genomes from the archaeal database used in the analysis can be found in Supplementary Table 6. For both phylogenies, the protein sequences for the analysis were aligned using clustalw v.2.1 with default settings ^105^. The aligned file was used to calculate a phylogenetic tree using RAxML v. 8.2.12 ^100^ with the following parameters “-m PROTGAMMAAUTO-f a-N 100-k”. The tree was then visualized using iTol^101^.

For the distribution and phylogenetic analysis of MCP and pPolB, known sequences encoded by various bacterial and archaeal viruses were used to build a Hidden Markov Model (HMM) via hmmer v3.3.2^106^. The HMM was then used to capture the corresponding components in proteomes of ANME-1 viruses and other MGEs. All sequences were then aligned using MAFFT v7.475^107^ option linsi and trimmed using trimAl v1.4.1^99^ option gappyout for pPolB and 20% gap removal option for MCP. Maximum-likelihood analyses were carried out through IQtree v2.1.12^108^ using model finder and ultrafast bootstrap with 2000 replicates. The phylogenetic tree was visualized and prepared using iTOL^101^.

For the distribution and phylogenetic analysis of ThyX, all ThyX sequences annotated by EggNOG mapper^109^ v2 in the genomes of ANME-1 and their MGEs were used to create a HMM as described above, and used to search for close homologs in the GTDB202 database, IMGVR V.3 database, as well as again in the proteomes and ANME-1 and their MGEs in this study. This yielded 261 sequences, which was then aligned and phylogenetically analyzed as for pPolB.

### CRISPR analysis

The CRISPR/Cas systems from the ANME-1 genomes and various metagenomic assemblies were annotated using CRISPRCasTyper v.1^49^. CRISPR spacer mapping onto MGEs was carried out as previously described^50^ with the following modifications. To filter out unreliable sequences that may have arisen during MAG binning, we took a conservative measure of only retaining CRISPR repeats identified in at least three ANME-1 contigs. We additionally analyzed the CRISPR repeats found in the *Alkanophagales* sister clade to ANME-1 using the same approach, which were found to have no overlap with the ANME-1 CRISPR repeats. To further avoid accidental mapping to unrelated MGEs, we applied a second stringent criteria of only retaining MGEs with at least 3 ANME-1 protospacers. MGEs larger than 10kb in size were retained for further analyses in this study.

### MGE network analysis and evaluation

Open reading frames in all CRISPR-mapped MGE contigs were identified using the PATRIC package^110^. Gene similarity network analyses were done using vCONTACT2 using the default reference (RefSeq202), with head-tailed viruses targeting haloarchaea and methanogens added as additional references^64^. Inverted and direct terminal repeats were detected using CheckV and the PATRIC package to determine genome completeness^110^.

### MGE annotation and virus identification

MGE proteomes are annotated using sensitive hidden Markov model profile-profile comparisons with HHsearch v3.3.0^111^ against the following publicly available databases: Pfam 33.1, Protein Data Bank (25/03/2021), CDD v3.18, PHROG (PMID: 34377978) and uniprot_sprot_vir70 (09/02/2021)^112^. Putative major capsid proteins of ‘Chaacviridae’ and ‘Ixchelviridae’ could not be identified using sequence similarity-based approaches. Thus, the candidate proteins were subjected to structural modeling using AlphaFold2^61,113^ and RoseTTAFold^6^. The obtained models were visualized using ChimeraX^114^ and compared to the reference structure of the major capsid protein of corticovirus PM2 (PDB id: 2vvf). The contigs containing identifiable viral structural proteins are described as viruses. The remaining contigs are described as unclassified MGEs, including circular elements that are most likely plasmids of ANME-1 as well as possible viruses enveloped by yet unknown structural proteins.

### Genome-scale virus comparisons

The viral genomes were annotated using Prokka v1.14.6^85^ to produce genbank files. Select genbank files were then analyzed using Clinker v. 0.0.23^115^ to produce the protein sequence clustering and alignments. Proteome-scale phylogeny for the head-tailed viruses were carried out via the VipTree server^65^.

### Taxonomic description of ‘*Ca*. Methanospirareceae’

Phylogenomic analysis placed the MAGs belonging to ANME-1c into two different genera, represented by one species in each (*‘Candidatus* Methanoxibalbensis ujae’ and ‘*Candidatus* Methanospirare jalkutatii’). Both genera belong to the same family named ‘*Candidatus* Methanospirareceae’ that it is included within the order ‘*Candidatus* Methanophagales’.

#### Description of the proposed genus ‘*Candidatus* Methanospirare jalkutatii’

(N.L. neut. n. *methanum* methane; N.L. pref. *methano-*, pertaining to methane; L. v. *spirare* to breathe; N.L. neut. n. *Methanospirare* methane-breathing organism; N.L. masc. n. *jalkutatii*, mythical dragon of the Pa ipai cosmology from Baja California, this dragon inhabited a beautiful place made of rocks and water similar to the Auka vent site). This organism is not cultured and is represented by five MAGs, all of them recovered from the hydrothermal environment of South Pescadero Basin. The type material is the genome designated FWG175, a single-scaffolded MAG comprising 1.99 Mbp in 1 circular scaffolds. This MAG was recovered from a methane-fed incubation of the mineral sample 12019 retrieved from the hydrothermal environment of South Pescadero Basin.

#### Description of the proposed genus ‘*Candidatus* Methanoxibalbensis ujae’

(N.L. neut. n. *methanum* methane; N.L. pref. *methano-*, pertaining to methane; N.L. adj. *xibalbensis* from the place called *Xibalba*, the Mayan word for the underworld; N.L. neut. n. *Methanoxibalbensis* methane-cycling organism present in deep-sea hydrothermal sediments; N.L. neut. adj. *ujae*, from the Kiliwa word *ujá* from Baja California meaning rock, referred to the high abundance of this species in rock samples).

This organism is not cultured and is represented by three MAGs from sedimented hydrothermal vents in the Gulf of California, one recovered from the Guaymas Basin and two from the South Pescadero Basin (see Supplementary Table 2). This group presumably is meso- or thermophilic and inhabits hydrothermal deep-sea environments, being mostly detected in rock samples. The type material is the genome designated NA091.008_bin2, a MAG comprising 1.96 Mbp in 86 scaffolds. The MAG was recovered from mineral sample (NA091.008) from the hydrothermal environment of South Pescadero Basin. It is proposed to be capable of anaerobic methanotrophy.

#### Description of the proposed family ‘*Candidatus* Methanospirareceae’

(N.L. neut. n. *Methanospirare* a (Candidatus) genus name; *-aceae* ending to denote a family). N.L. neut. pl. n. *Methanospirareceae* the (Candidatus) family including the (Candidatus) genera of *Methanoxibalbensis* and *Methanospirare*. The description is the same as for the candidate type genus *Methanospirare*. The 16S rRNA genes of this family correspond to the previously described 16S rRNA gene sequences known as “ANME-1 Guaymas”^18^.

### Taxonomic description of proposed ANME-1 virus orders and families with representative complete genomes

#### Order *Coyopavirales*, family *Chaacviridae*

This group of viruses is characterized by novel major capsid protein (MCP), which is predicted using AlphaFold2 to have the double jelly-roll (DJR) fold, the hallmark protein of viruses within the realm *Varidnaviria*. We propose classifying these viruses into a new family, *Chaacviridae*, after Chaac, the god of death in the Mayan mythology. Chaacviruses displays minimal proteome overlap with other known viruses, and is characterized by a uniform 10-11 kb genome size and a gene encoding protein-primed family B DNA polymerase (pPolB).

*Chaacviridae* comprises two genera that have relatively similar functional composition in their proteomes, yet exhibit relatively low sequence conservation and distinct gene arrangements. Viruses in the two genera appeared to have undergone a genomic inversion during their evolutionary history. We propose the genus names *Homochaacvirus* and *Antichaacvirus* (from *homo* [same in Greek] and *anti* [opposed in Greek] to emphasize the inversion of a gene module including the *pPolB* gene). Three complete genomes of chaacviruses have been obtained, as judged from the presence of inverted terminal repeats, consistent with the presence of *pPolB* gene. The three viruses share less than 90% average nucleotide identity and represent separate species, respectively named *Homochaacvirus pescaderoense* PBV0304, *Homochaacvirus californiaense* PBV0305, and *Antichaacvirus pescaderoense* PBV266.

While *Chaacviridae* is not closely related to any of the existing virus families, based on the gene complement, chaacviruses resemble members of the class *Tectiliviricetes*. Thus, we propose placing *Chaacviridae* into a new monotypic order *Coyopavirales* (after Coyopa, the god of thunder in Mayan mythology) within the existing class *Tectiliviricetes*.

#### Order *Nakonvirales*, families *Ahpuchviridae* and *Ekchuahviridae*

Based on ViPTree analysis, all ANME-1 viruses belonging to *Caudoviricetes* form a distinct clade outside of the three existing orders of haloarchaeal and methanogenic archaeal viruses ^64^ (Fig. 4). We propose to create a new order *Nakonvirales* (after Nakon, the most powerful god of war in Mayan mythology) for unification of two family-level groups that have complete genome representatives.

We propose naming the first of the two groups *Ahpuchviridae* (after Ah Puch, the god of death in the Mayan mythology). This family is represented by one genus *Kisinvirus* (after Kisin, another Mayan god of death) and a single species, *Kisinvirus pescaderoense*. The species includes virus PBV299, which has a dsDNA genome of 70,925 bp and besides the morphogenetic genes typical of members of the *Caudoviricetes*, encodes an RNA-primed family B DNA polymerase, archaeo-eukaryotic primase and a processivity factor PCNA.

The second proposed family, *Ekchuahviridae* (after Ek Chuah, the patron god of warriors and merchants in the Mayan mythology), is represented by one genus *Kukulkanvirus* (after Kukulkan, the War Serpent in the Mayan mythology). The proposed genus will include two species, *Kukulkanvirus guaymasense* and *Kukulkanvirus mexicoense*, with representative viruses containing genomes of 71,795 bp and 80,551 bp, respectively. Notably, viruses in this group encode two divergent HK97-fold MCPs with their own capsid maturation proteases, but all other canonical head-tailed virus structural proteins are encoded as single copy genes. Their replication modules include RNA-primed family B DNA polymerase and archaeo-eukaryotic primase.

#### Order *Maximonvirales*, family *Ahmunviridae*

For classification of rod-shaped virus PBV300, we propose the creation of a new genus, *Yumkaaxvirus* (after Yum Kaax, the god of the woods, the wild nature, and the hunt in Mayan mythology) within a new family, *Ahmunviridae* (after Ah Mun, the god of agriculture in Mayan mythology). PBV300 has a linear dsDNA genome of 41,525 bp with 99 bp terminal inverted repeats. It encodes two divergent MCPs homologous to those of viruses in the realm *Adnaviria*. The virus also encodes several other proteins with homologs in members of the family *Rudiviridae*, including the terminal fiber protein responsible for receptor binding. This family is related to members of the class *Tokiviricetes* (realm *Adnaviria*), but outside of the two existing orders, *Ligamenvirales* and *Primavirales*. We thus propose to assign *Ahmunviridae* to a new order *Maximonvirales*, after Maximon, a god of travelers, merchants, medicine men/women, mischief and fertility in Mayan mythology.

#### Family *Itzamnaviridae*

We propose the family name *Itzamnaviridae* (after Itzamna, lord of the heavens as well as night and day in the Mayan mythology) for the spindle-shaped viruses with complete genomes in this study. The members of this family differ in genome sizes and are subdivided into two genera, which we propose naming *Demiitzamnavirus* and *Pletoitzamnavirus* (after *demi-* for half or partial [derived via French from Latin ‘*dimedius’]* and *pleto* for full [Latin]). Demiitzamnaviruses have circular genomes sized around 25 kb, each encoding two MCPs homologous to those characteristics of and exclusive to archaeal spindle-shaped viruses. This genus is represented by *Demiitzamnavirus mexicoense* GBV303. Pletoitzamnaviruses have genome sizes around 45-48 kb, where roughly half of the genome aligns with nearly the entirety of the genomes of demiitzamnaviruses, including the two MCP genes. However, the remaining fraction of pletoitzamnavirus genomes encodes enzymes of diverse functions, including replicative proteins, such as RNA-primed family B DNA polymerase and archaeo-eukaryotic primase, PCNA, large and small subunit of the replication factor C, etc. The content of this genome fraction is relatively flexible, with genes encoding various metabolic and regulatory enzymes swapping in and out, often in the form of multi-gene cassettes. For example, the representative *Pletoitzamnavirus pescaderoense* PBV082, contains a gene cluster encoding a kinase-phosphatase pair, a thymidylate synthase, and a radical SAM enzyme of unknown function. Except for the characteristic MCP, members of the *Itzamnaviridae* do not encode proteins with appreciable sequence similarity to proteins of other spindle-shaped viruses.

A formal proposal for classification of the ANME-1 viruses discovered in this study has been submitted for consideration by the International Committee for Taxonomy of Viruses (ICTV) and are detailed in Supplementary Table 12.

## Data availability

Raw metagenome reads, assembled metagenome bins and virus sequence data are available in GenBank under BioProject accession numbers PRJNA875076 and PRJNA721962. Select complete ANME-1 virus genomes can be found on GenBank under accession numbers OP413838, OP413839, OP413840, OP413841, OP548099, and OP548100.

## Acknowledgements

We are indebted to the crews from R/V Falkor (cruise FK181031) and E/V Nautilus (cruise NA091) and the pilots of ROVs SuBastian and Hercules. Sample collection permits for FK181031 (25/07/2018) were granted by la Dirección General de Ordenamiento Pesquero y Acuícola, Comisión Nacional de Acuacultura y Pesca (CONAPESCA: Permiso de Pesca de Fomento No. PPFE/DGOPA-200/18) and la Dirección General de Geografía y Medio Ambiente, Instituto Nacional de Estadística y Geografía (INEGI: Autorización EG0122018), with the associated Diplomatic Note number 18-2083 (CTC/07345/18) from la Secretaría de Relaciones Exteriores - Agencia Mexicana de Cooperación Internacional para el Desarrollo / Dirección General de Cooperación Técnica y Científica. Sample collection permit for cruise NA091 (18/04/2017) was obtained by the Ocean Exploration Trust under permit number EG0072017. This research used samples provided by the Ocean Exploration Trust’s Nautilus Exploration Program, cruise NA091. E/V Nautlius operated by the Ocean Exploration Trust, with cruise NA091 supported by the Dalio Foundation and Woods Hole Oceanographic Institute, and R/V Falkor operated by the Schmidt Ocean Institute. Funding for this work was provided by grants from the National Science Foundation Center For Dark Energy Biosphere Investigations (C-DEBI) and the NOMIS foundation (V.J.O.). VJO contribution was supported by the U.S. Department of Energy, Office of Science, Office of Biological and Environmental Research under Award Number DE-SC0020373. This work was also supported by the Deutsche Forschungsgemeinschaft (DFG, German Research Foundation) under Germany’s Excellence Initiative/Strategy through the Cluster of Excellence “The Ocean Floor-Earth’s Uncharted Interface” (EXC-2077-390741603 R.L.-P. and V.J.O). M.K. was supported by l’Agence Nationale de la Recherche grant ANR-20-CE20-0009-02. F.W. was supported by the Dutch Research Council Rubicon Award 019.162LW.037, the Human Frontiers Science Program Long-term fellowship LT000468/2017, and a ZJU-HIC Independent PI Startup Grant.

## Disclaimer

This report was prepared as an account of work sponsored by an agency of the United States Government. Neither the United States Government nor any agency thereof, nor any of their employees, makes any warranty, express or implied, or assumes any legal liability or responsibility for the accuracy, completeness, or usefulness of any information, apparatus, product, or process disclosed, or represents that its use would not infringe privately owned rights. Reference herein to any specific commercial product, process, or service by trade name, trademark, manufacturer, or otherwise does not necessarily constitute or imply its endorsement, recommendation, or favoring by the United States Government or any agency thereof. The views and opinions of authors expressed herein do not necessarily state or reflect those of the United States Government or any agency thereof.

## Author contributions

R. L.-P., F.W., A.C., and V.J.O. conceived and designed the study. D.R.S, V.J.O. and J.S.M. retrieved the original samples. A.C. and F.W. carried out rock incubations and FISH microscopy. R.L.-P., F.W. and A.C. performed DNA extraction. R.L.-P. and F.W. performed metagenomic assembly and analysis. F.W. performed CRISPR-based mobilome discovery. M.K. and F.W. performed analyses of viruses. R.L.-P., F.W., M.K. and V.J.O wrote the manuscript with contributions from all coauthors. We declare no competing financial interests.

---

**Supplemental Figure 1.**
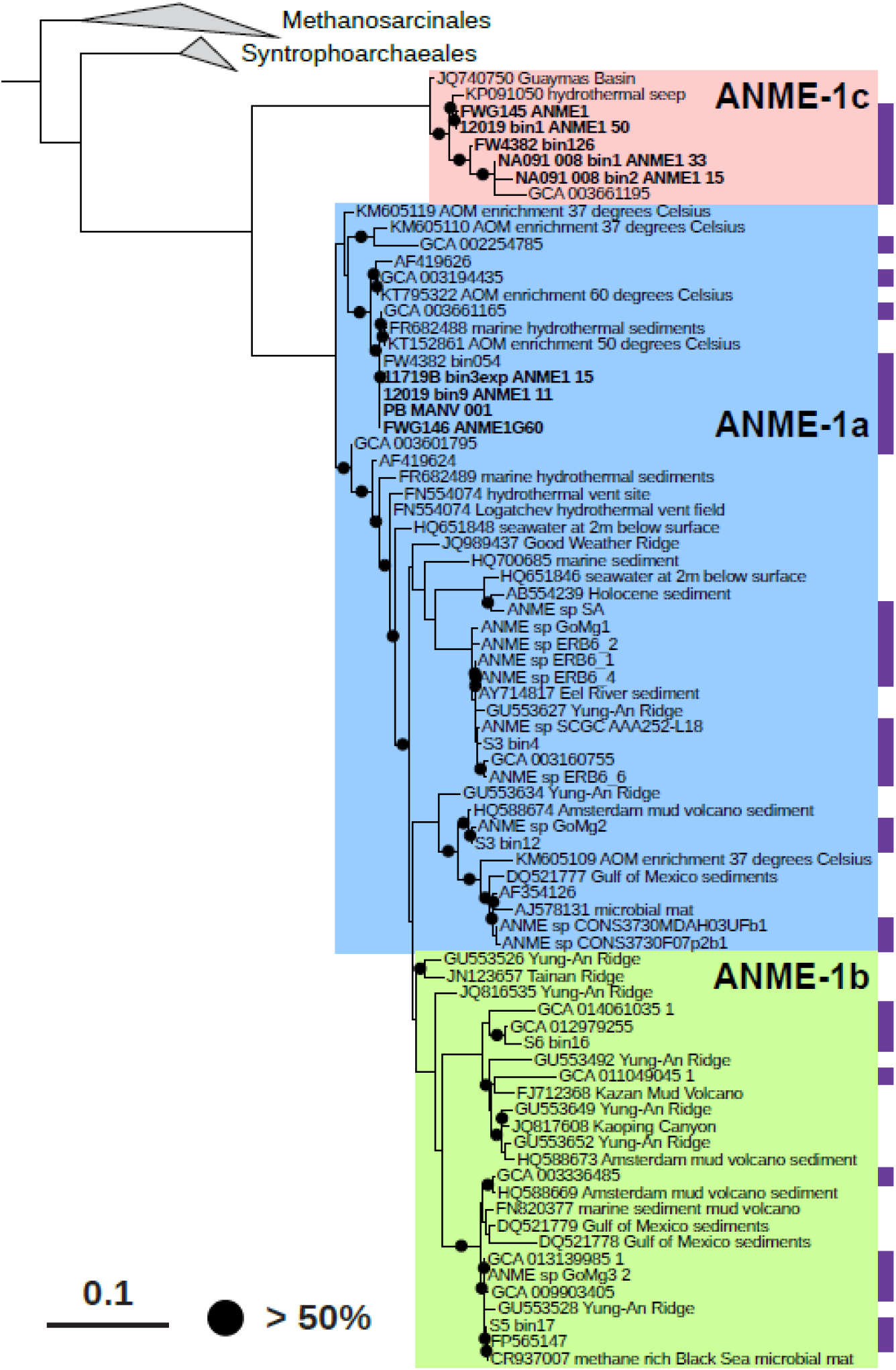
16S rRNA gene phylogeny for the ANME-1 clade (*Methanophagales*). Color shading highlights the three main groups of ANME-1 archaea. The purple bars note 16S rRNA gene sequences retrieved from MAGs shown in Figure 1. Sequences retrieved from Pescadero MAGs are in bold. Bootstrap values over 50% are indicated with a black circle. Scale bar indicates the number of nucleotide substitutions per site.

**Supplemental Figure 2.**
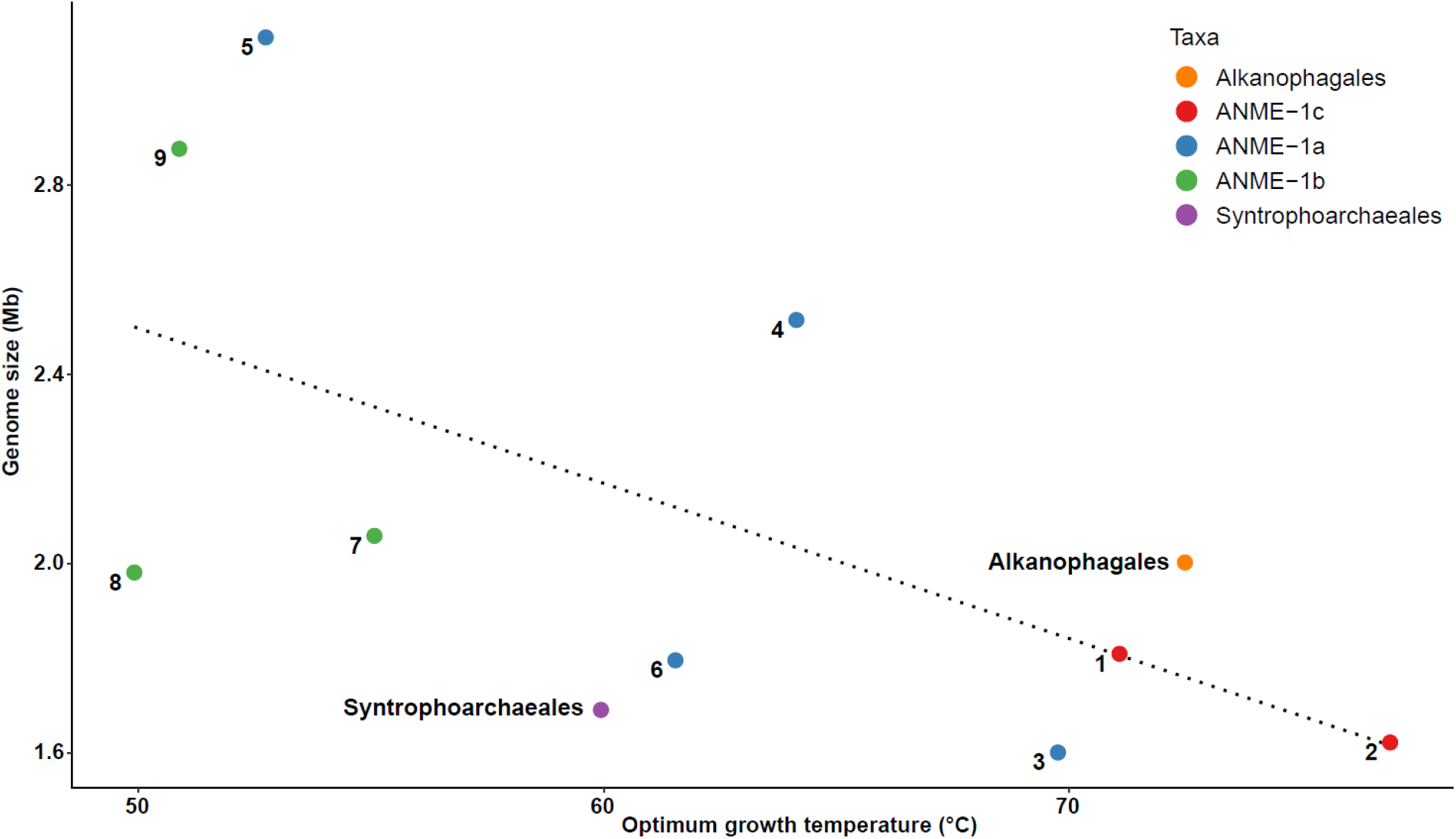
Correlation between estimated genome size (in Mb and after calculation considering contamination and completeness see Material and Methods) and the predicted optimum growth temperature (°C). Each point and number represents the average values for one ANME genera/species (see Supplementary Table 2), except in the case of Syntrophoarchaeales and Alkanophagales where the point represent the average values for the whole clade. Color indicates the corresponding taxonomy. Dotted line indicates the regression model (R^2^=0.3858).

**Supplemental Figure 3.**
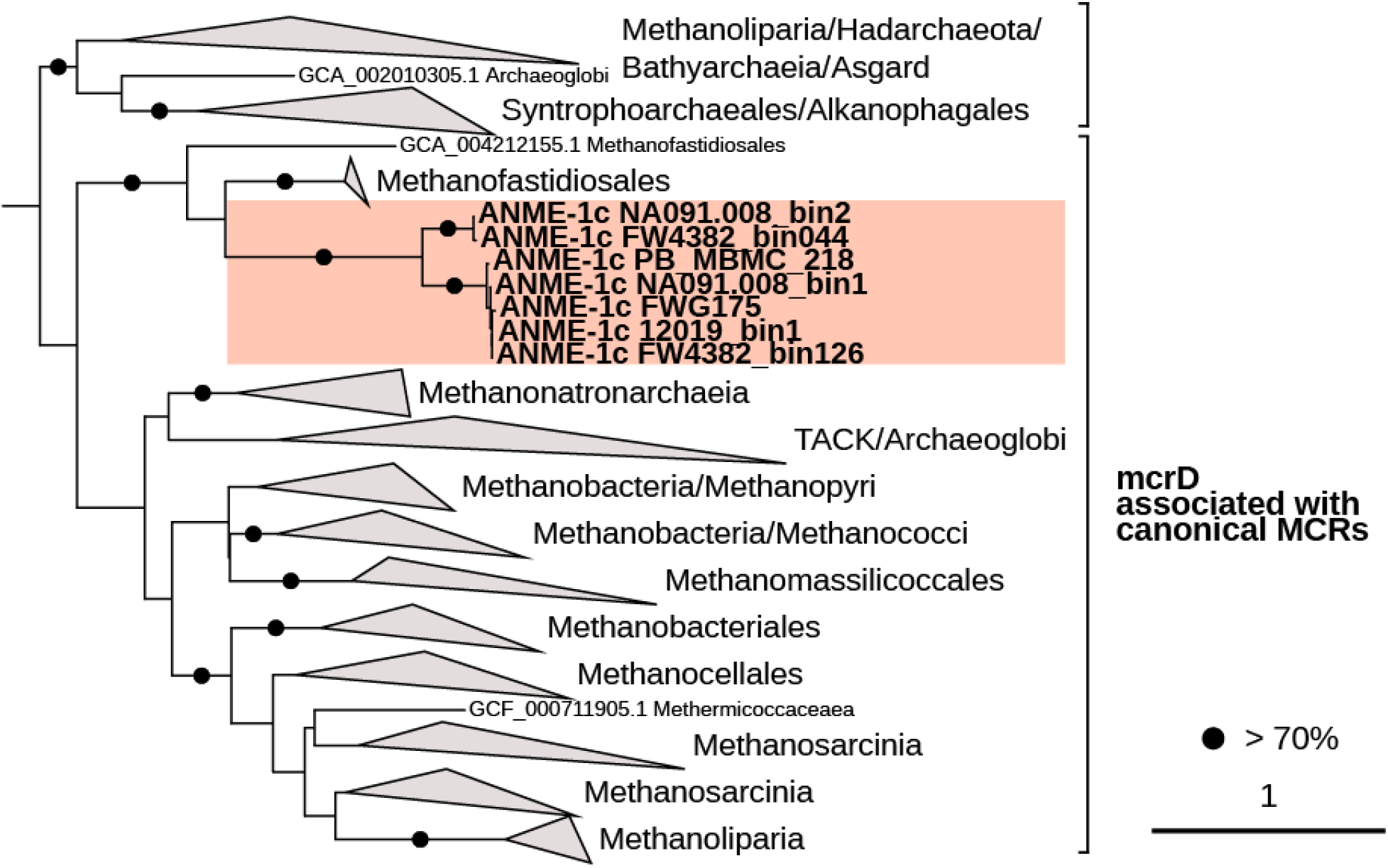
Phylogenetic tree of McrD genes from archaea, including the McrD in ANME-1 genomes (only found in ANME-1c). Black circles indicate bootstrap support values over 70%. Scale bar represents the number of amino acid substitutions per site.

**Supplemental Figure 4.**
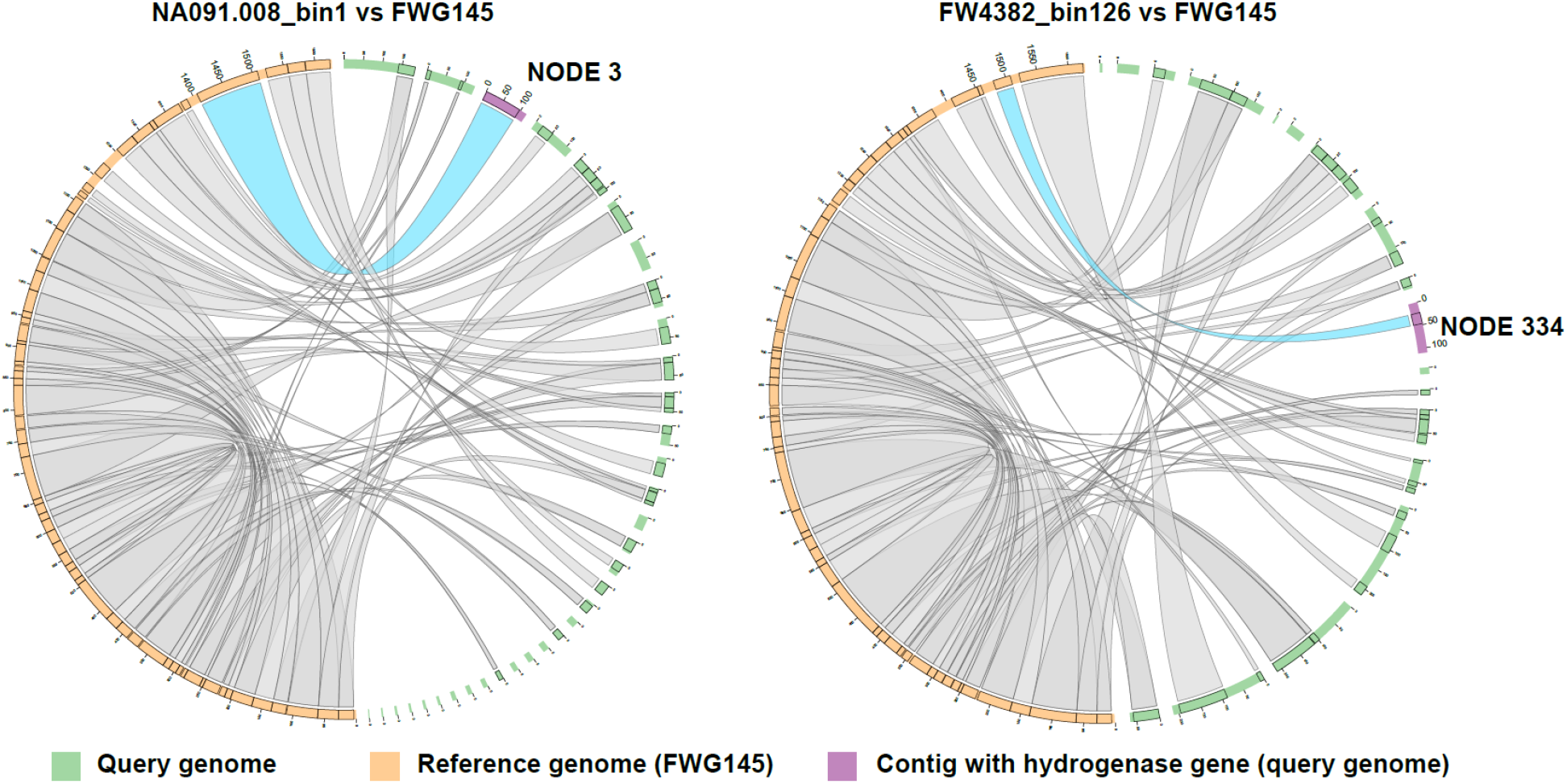
Circos plot comparing homologous regions of the ANME-1c genomes, NA091.008_bin1 and FW4382_bin126 (both with hydrogenase operons) to the predicted completed genome FWG175 that was assembled as a single contiguous scaffold and belongs to the same species. Contigs corresponding to the query genomes (NA091.008, FW4382_bin126) are marked in green and contigs from genome FWG175 are in orange. The contig containing the hydrogenase operon is shown in purple and the corresponding homology sections between the reference and query genomes are highlighted in blue. The region between these homology sections corresponds to the hydrogenase operon that was not detected in genome FWG145.

**Supplemental Figure 5.**
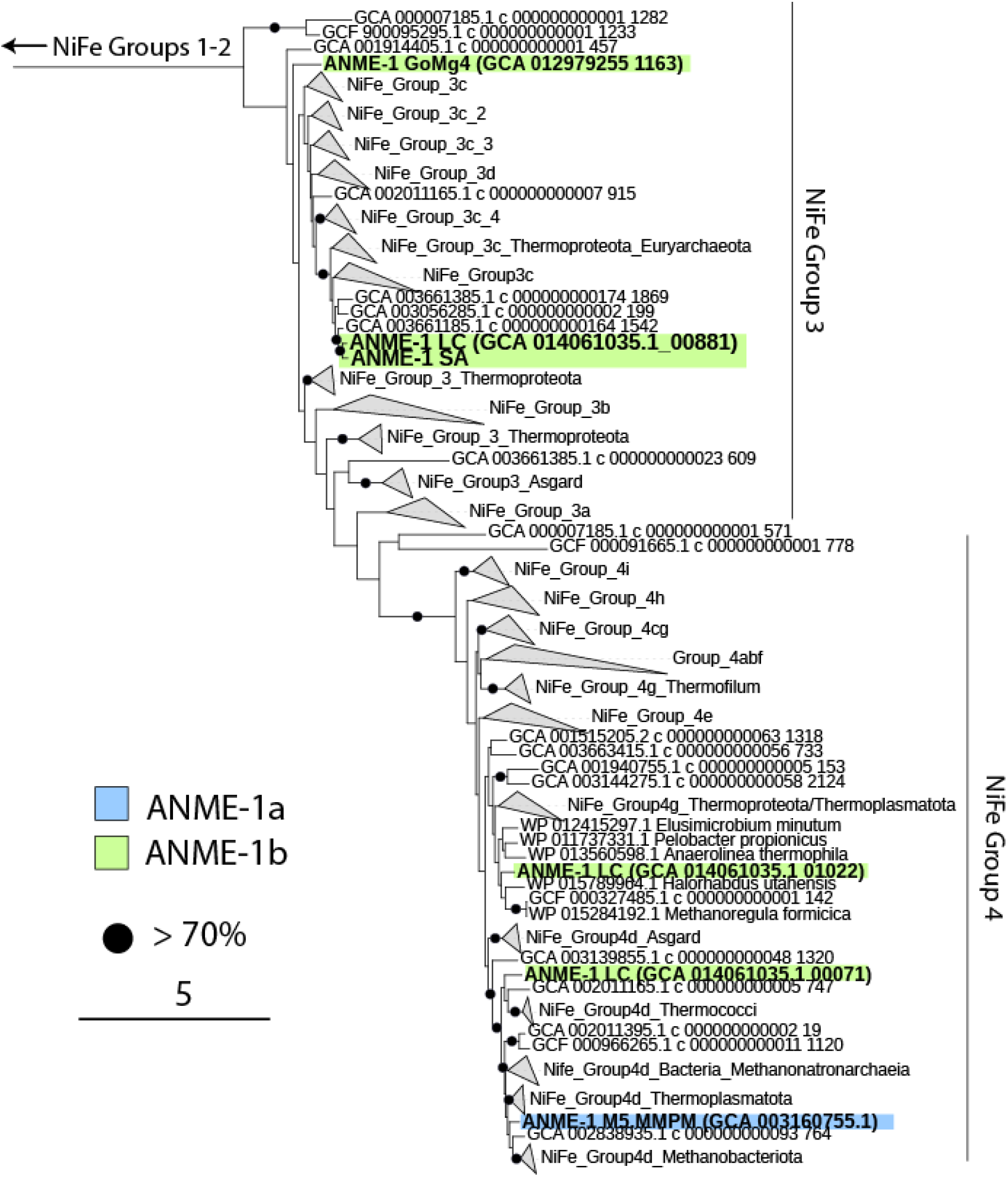
Phylogenetic tree of the large subunit of the NiFe hydrogenase present in ANME-1 genomes associated with NiFe Groups 3 and 4. The green and blue shading indicates the taxonomic identity of the ANME-1 MAG containing the corresponding hydrogenase. Black circles indicate bootstrap support values over 70% (left). The scale bar represents the number of amino acid substitutions per site.

**Supplemental Figure 6.**
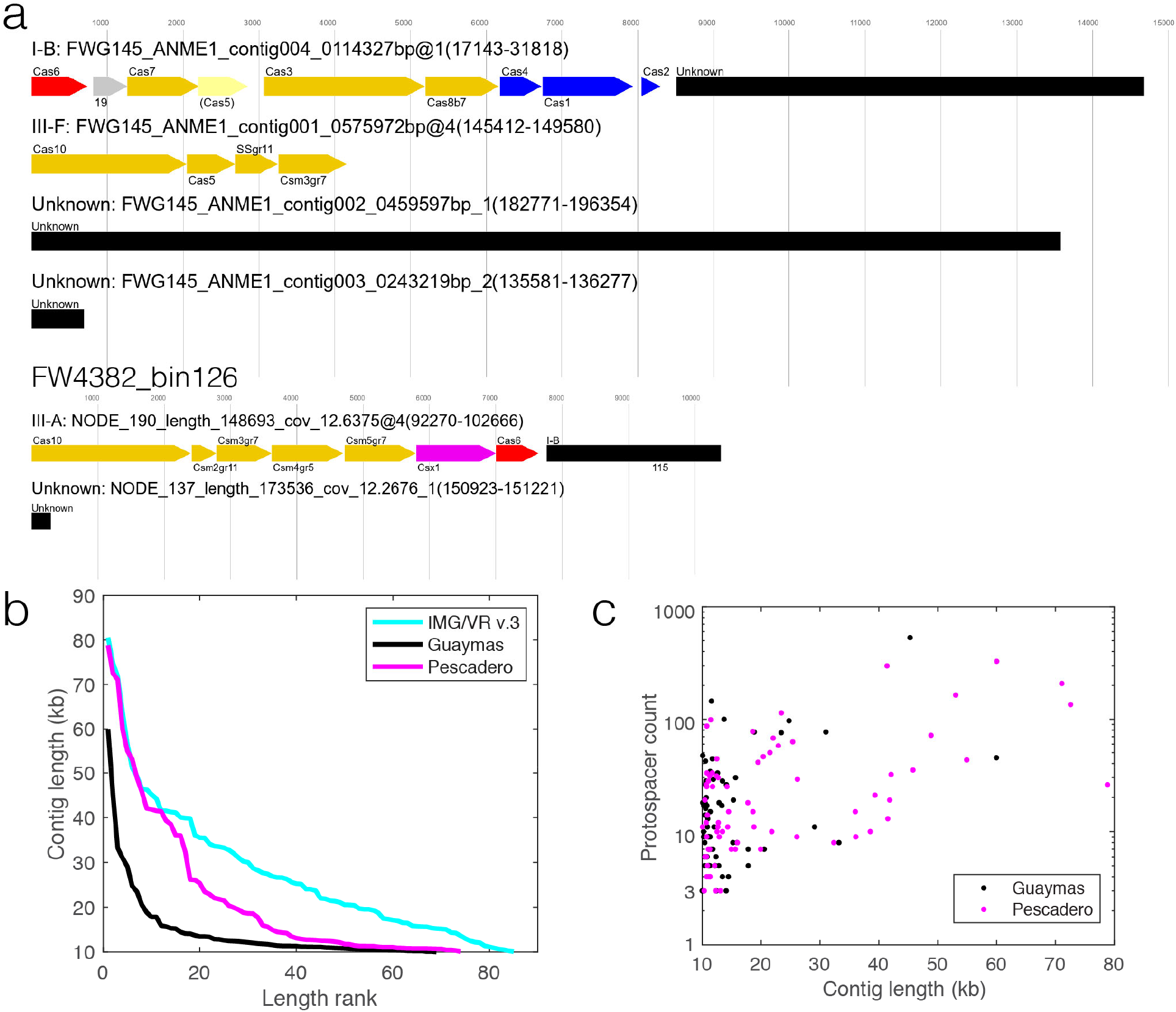
Features of ANME-1 CRISPR/Cas and spacer-mobilome mapping. **a**) CRISPR/Cas features in the two most contiguous ANME-1c MAGs characterized using CCtyper. Black bars indicate CRISPR arrays. **b**) Contig lengths of all ANME-1 mobile genetic elements (MGEs) found in this study. Note that the distribution does not necessarily indicate completeness as IMG/VR v.3 is more enriched in head-tailed viruses (with genomes sized up 80 kb) whereas the Pescadero/Guaymas basin dataset contains many tailless icosahedral viruses whose genomes are sized around 10 kb. **c**) Distribution of protospacers within the ANME-1 mobile elements found in S. Pescadero and Guaymas basins.

**Supplemental Figure 7.**
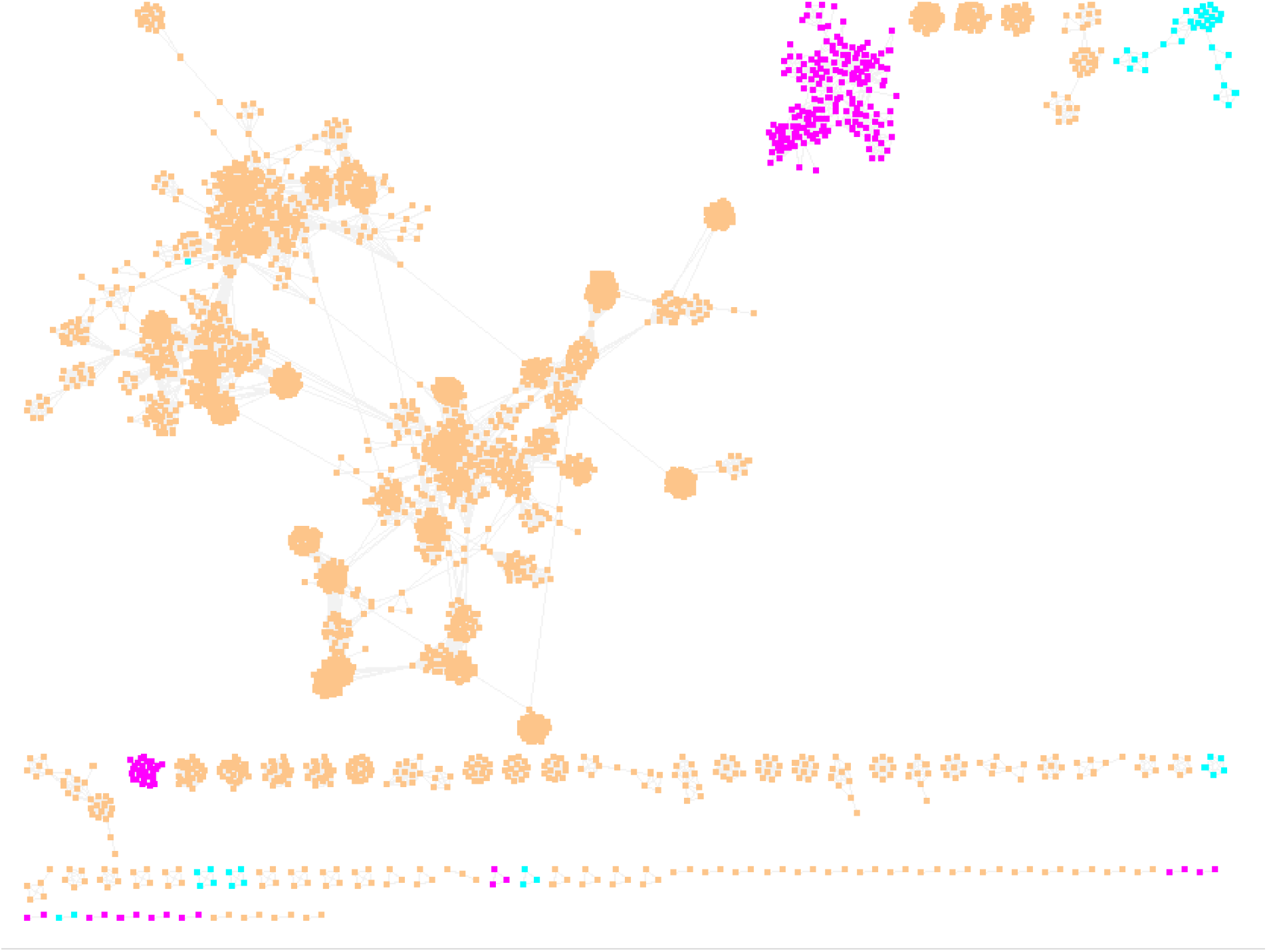
Gene-sharing networks produced via vCONTACT2 indicate that all ANME-1 mobile genetic elements (Magenta) are well distinguished from the known *haloarchaeal* viruses, or *Haloviruses*, (Blue) and other viruses with known hosts (orange).

**Supplemental Figure 8.**
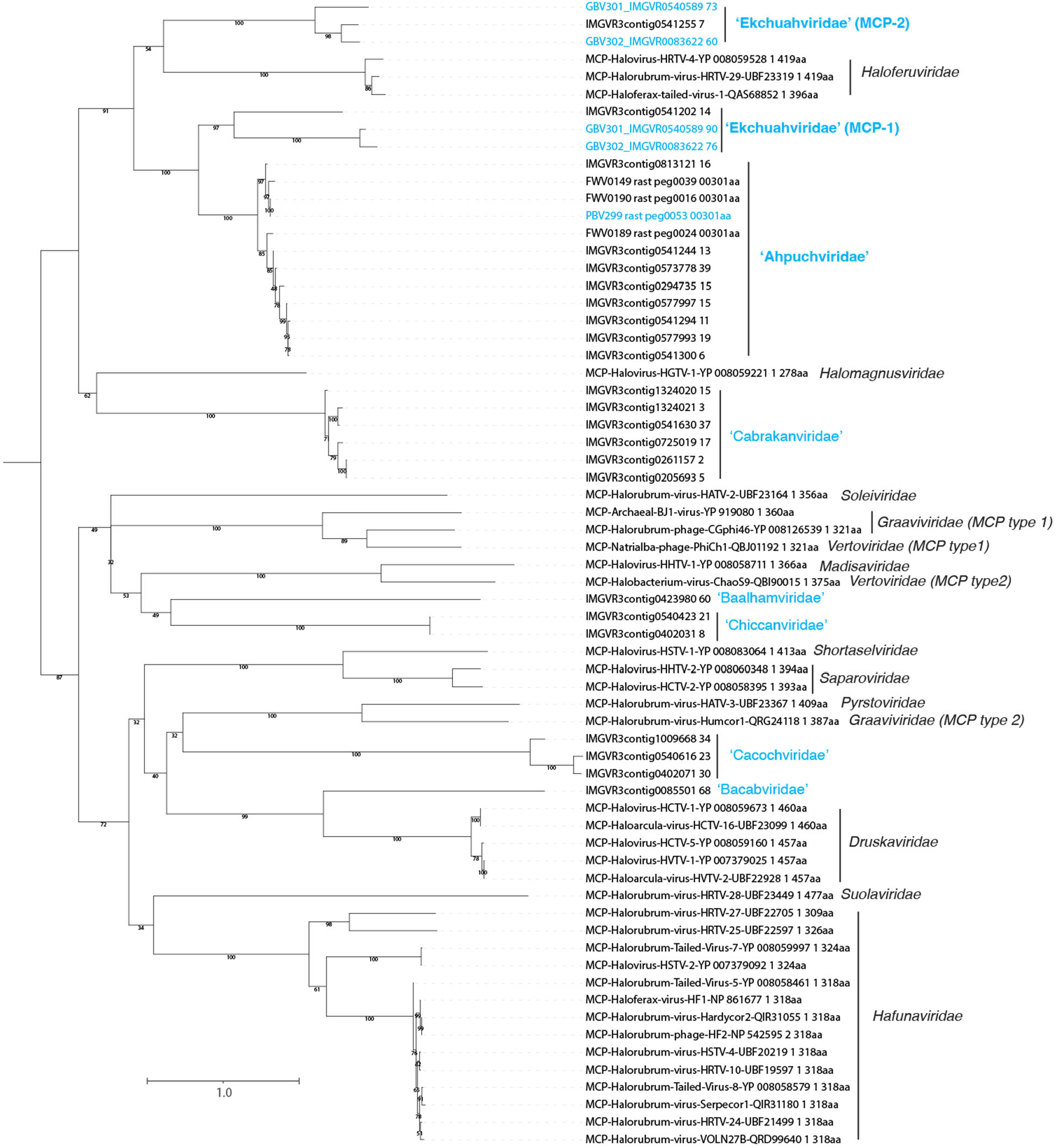
Maximum likelihood analyses of MCPs encoded by head-tailed viruses. Blue, ANME-1 virus families; black, haloarchaeal virus families. MCPs from complete genomes of ANME-1 viruses are indicated in blue, and their respective families in bold.

**Supplemental Figure 9.**
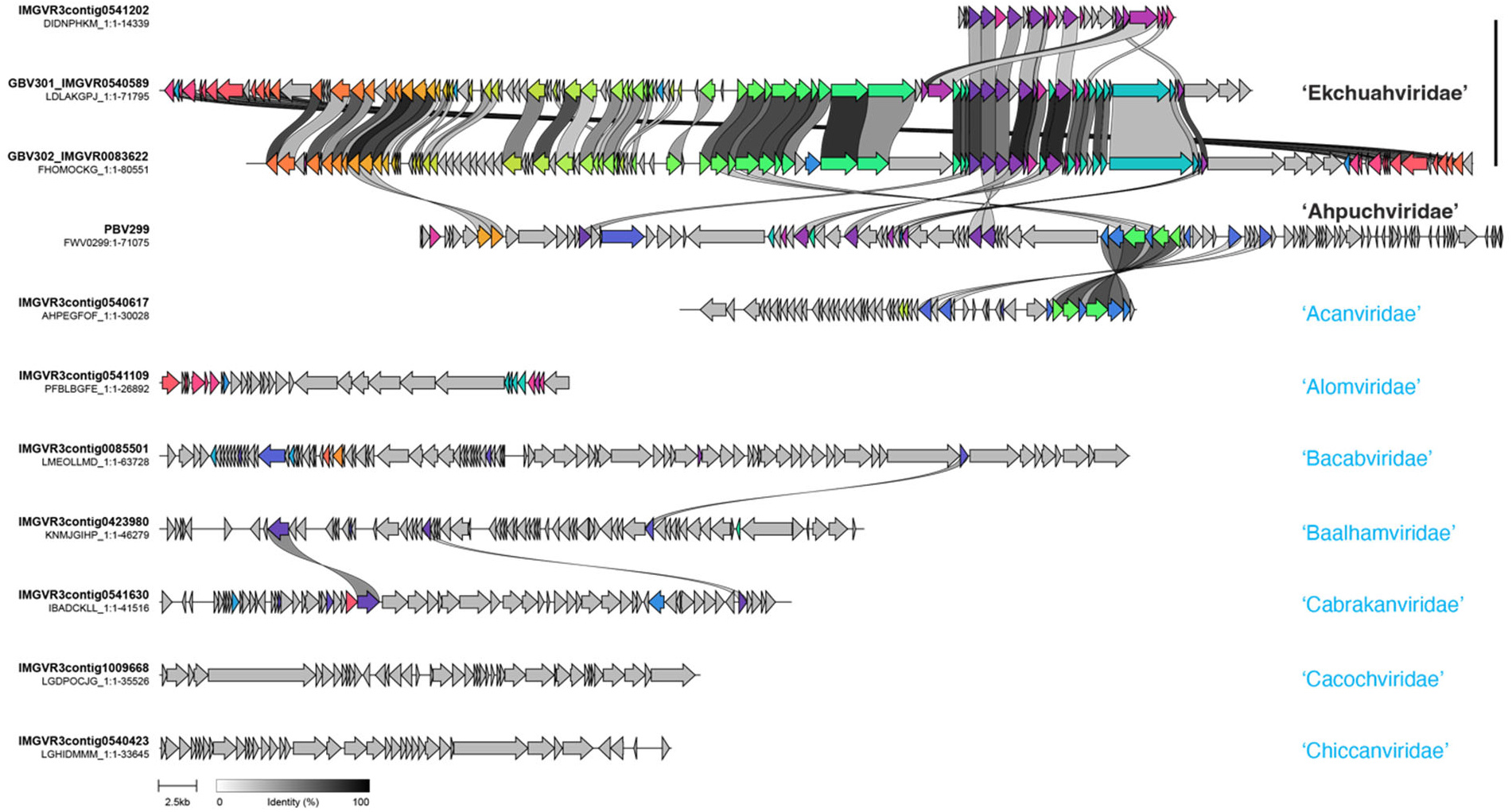
Sequence alignment of representatives of ANME-1 viruses. Colors indicates protein families encoded by at least 2 representative viral genomes here. Grey indicates singleton proteins without apparent homologs. Scale bars for protein identity scores and genome sizes are indicated at the bottom. Viral contig names and sizes are indicated on the left side and their respective family and Candidatus (Ca.) family names are indicated on the right.

**Supplemental Figure 10.**
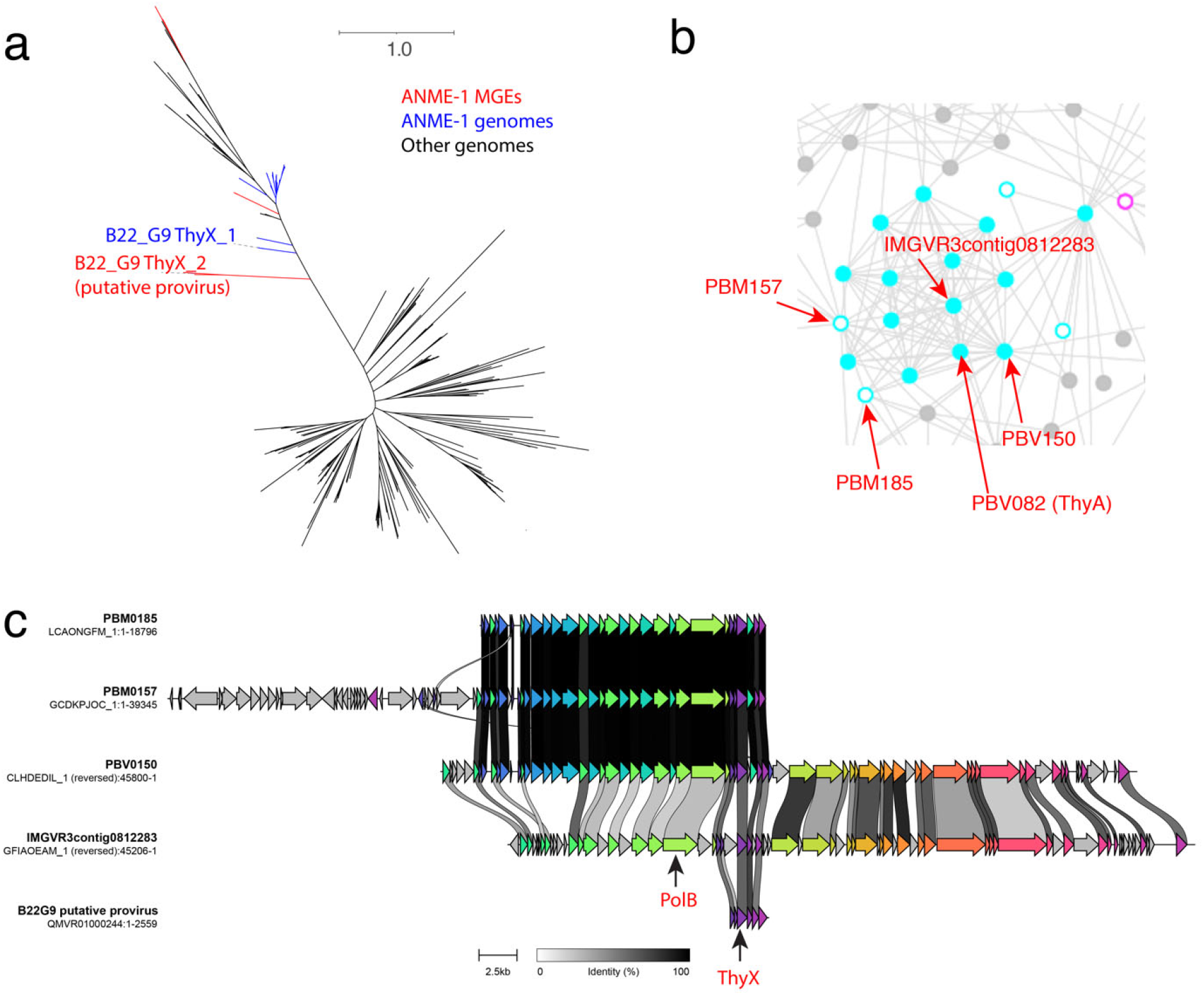
Spindle-shaped viruses encode ThyX at the root of ANME-1 ThyX. a, unrooted phylogeny suggests that ANME-1 *thyX* may have evolved from *thyX* genes originated in ANME-1 viruses. The two versions of of ThyX in the ANME-1c bin B22_G9 is highlighted. b, two MGEs containing thyX genes are found to be highly related to the spindle-shaped viruses with identified MCPs. c, Sequence alignment indicate that the two MGEs and a genome contig that encode ThyX are all fragmented from spindle-shaped viruses.

