## Supplementary Text for "Evolutionary Diversification of Methanotrophic *Ca*. Methanophagales (ANME-1) and Their Expansive Virome"

Coenzyme M biosynthesis

Coenzyme M is a fundamental coenzyme for methanogens and ANME organisms. So far, two different routes (I and II) for CoM biosynthesis are described in methanogens[1-3]. The pathway I appears in Class I methanogens (*Methanococcales, Methanobacteriales* and *Methanopyrales*), while pathway II is present in Class II methanogens like *Methanosarcinales* and *Methanomicrobiales*[2]. Both synthetize CoM from 3-sulfopyruvate and they shared the enzyme sulfopyruvate decarboxylase, either as a monomeric enzyme (comDE) or as a dimeric form (comD+E)[4]. Presumably, both routes also share the recently described CoM synthase (comF), that catalyze the last step of the route: the conversion of sulfoacetaldehyde to CoM[3]. The routes differ on the way to produce 3-sulfopyruvate. In pathway I sulfite and phosphoenolpyruvate are combined to form 3-sulfopyruvate in a multi-step process catalyzed by three different enzymes (comA, comB, comC)[2], whereas in the pathway II this molecule is produced from *O*-phospho-L-serine in two steps via L-cysteate and cysteate aminotransferease.

CoM biosynthesis does not seem to be a conserved trait in ANME-1. Both pathways are present in different ANME-1 groups (Figure 1). ANME-1c, two ANME-1a clades and one ANME-1b group seem to use the biosynthesis pathway II since they encode for a cysteate synthase (MA_3297), a monomeric comDE and a comF. In these MAGs, we could not find genes for comA, comB and comC. We also were unable to identify a phosphoserine aminotransferase present in pathway II, although there were several uncharacterized genes with aminotransferase motifs. On the other side, two ANME-1a clades and an additional ANME-1b group seem to use the pathway I for CoM biosynthesis since they have genes for comA, comB, comF and a monomeric comDE. In this case, some genomes had genes for comC and some lacked them. This distribution suggest that the CoM biosynthesis pathway is not a monophyletic trait in ANME-1, although it is central route in ANME-1 metabolism. The sister clade of ANME-1, *Syntrophoarchaeales*, seem to utilize the biosynthetic pathway I since their MAGs encode for a comB, although we could not detect genes for comA and comC. The presence of the biosynthetic pathway II in some ANME-1 clades (including ANME-1c) indicates that the CoM biosynthesis was likely acquired at different moments and possibly by different clades.

Archaellum and chemotaxis system

It was particularly intriguing to observe the presence of genes coding for an archaellum, an archaeal motility structure, and an elaborate chemotaxis system in some of the MAGs, including most of the ANME-1c. The archaellum consists of several proteins (Arl) like structural pilins known as archaellins (FlaB, FlgA or ArlA), a peptidase to process the pre-archaellins (FlaK or ArlK), an ATPase for the assembly of the pilin subunits (ArlI or FlaI) and different membrane proteins with a motor or switch function (ArlH, ArlCDE, ArlJ or FlaH, FlaCDE, FlaJ) [5 6]. In endolithic habitats inhabited by ANME-1c, the archaellum might give a selective advantage in different ways: conferring cellular motility[5], as attachment element[7,8] or acting as a conductive nano-wire as has been shown in other archaea[9,10]. In porous or fractured rock matrices, these components might allow ANME-1c to search for the optimal environmental conditions within their heterogeneous and fluctuating habitats. Notably, Type IV pili, which function as receptors for diverse archaeal viruses[11,12], could mediate adhesion of ANME-1c to different surfaces

The prokaryotic chemotaxis system allows organisms to regulate the cell motility depending on environmental stimuli[13,14]. Several chemotaxis proteins are encoded across different ANME-1 groups, including some without an archaellum (Figure 1). For instance, genes for methyl-accepting chemotaxis proteins (MCP), the histidine kinase CheA, the adaptor protein CheW and the response regulators CheY and CheB are present in MAGs of the ANME-1c, ANME-1a and ANME-1b. Interestingly, only a few MAGs from ANME-1c and two Pescadero ANME-1a MAGs have three additional genes for chemotaxis: CheC, CheD and CheF. The latter play a fundamental role by transmiting the signal from the chemotaxis system to the archaellum via the FlaCDE proteins[15]. In the ANME-1c, these three genes appear only in MAGs from species 1, which possess an archaellum, while strikingly some genomes from species 2 have the machinery for an archaellum, but lack most of the chemotaxis system. Interestingly, some crenarchaea have a canonical archaellum, but they lack the traditional chemotaxis system[6], what led to hypothesize if uncharacterized chemotaxis systems exist in archaea[14]. The existence of chemotaxis genes in other ANME-1 that do not encode for an archaellum has still no explanation, but it may be related to the presence of diverse genes for type IV pili, which are widespread in ANME-1.

Nitrogen metabolism

Like many ANME-1 genomes (Figure 1), ANME-1c have genes encoding for some of the subunits of the nitrogenase enzyme, necessary for nitrogen fixation, like NifH and NifD, but lacks other genes for the rest of the subunits (NifK, $NifI_1$ and $NifI_2$). Therefore, there are doubts about the ability of ANME-1 to fix $N_2$[16,17]. Instead, the NifH and NifD present in ANME-1 are proposed to catalyze some steps in the biosynthesis of the $F_{430}$ cofactor[18]. ANME-1c might assimilate inorganic nitrogen as ammonium through a glutamine synthetase (glnA) as it was proposed for other ANME-1[16].

ANME-1c have several genes for the synthesis of polyamines (Figure 1), a group with a variety of organic molecules with at least two amino groups that perform diverse functions in archaea and bacteria[19]. Agmatine, putrescine and spermidine are some of the most studied polyamines. The ANME-1 order seem to be able to produce agmatine and putrescine, thanks to the genes encoding for arginine decarboxylase (speA) and agmatinase (speB). Additionally, the ANME-1c group a gene for a spermidine synthase (speE), which catalyze the production of spermidine, whose function in ANME-1c is still unknown. This gene is absent in most other ANME-1 genomes. In *Thermococcus kodakarensis*, spermidine is related to the production of longer- and brached-chain polyamines with a critical role for growth at high temperatures [20]. Therefore, the production of spermidine in ANME-1c may allow them to thrive at high temperatures, considering that the predicted optimal growth temperature of this group is over 70 °C and the highest of the whole ANME-1 order (Supplementary Table 4).

### Aminoacid metabolism

ANME-1c have the potential to synthetize most amino acids similarly to other ANME-1 genomes [17]. The only notable exception is the absence of some genes for the metabolism of proline and alanine that are present in other ANME-1 clades (Figure 1).

We could not detect two genes central in the biosynthesis of proline in any MAG of ANME-1c. Their products are a glutamate 5-kinase (proB) and a glutamate 5-semialdehyde dehydrogenase (proA), responsible for the conversion of glutamate to glutamate 5-semialdehyde[21]. We do detect a gene for pyrroline-5-carboxylate reductase (proC) that catalyze the last step in proline biosynthesis. Most likely ANME-1c possess an alternative route to replace the steps catalyzed by proB and proA.

Finally, ANME-1c appear to lack any homologues for the biosynthesis of alanine in ANME-1c, although they are present in other ANME-1 genomes. For instance, ANME-1 genomes usually have genes for the synthesis of alanine from pyruvate or valine or possess a copy for an L-cysteine desulfurase (csdA) that synthesize alanine from L-cysteine[17] and is involved in the biosynthesis of Fe-S centers [22]. . We could not identify any of them in ANME-1c. However, many of the biosynthetic enzymes for alanine possess the aminotransferase COG motif AspB (COG0436)[21], motif that was present in several of the encoded enzymes of ANME-1c. Moreover, *csdA* might be replaced in ANME-1c by a cysteine desulfidase (*cdsB*), which was suggested to be involved in the biosynthesis of Fe-S centers in the archaeon *Methanocaldococcus jannaschii*[23].

### Other hydrogenases in ANME-1

From the 33 hydrogenases detected in ANME-1, 26 fall in a monophyletic group next to the hydrogenases of the groups 1j and 1h. Only seven hydrogenases fall out of this group. One belongs to the ANME-1b SA genome and is affiliated to euryarchaeotal hydrogenases from *Archaeoglobi* and *Halobacteria*, between the NiFe Group 1j (*Archaeoglobi*, anaerobic) and the clade with the NiFe groups 1g, 1h and the rest of ANME-1 hydrogenases (Figure 3). Three hydrogenases (ANME-1 SA, ANME-1 GoMg4 and ANME-1 LC) are affiliated to hydrogenases of the group 3c, present in many methanogens, where they likely bifurcate electrons from $H_2$ to heterodisulfide and ferredoxin. These three ANME-1b species belong to the group *Methanoalium*, which is proposed to perform methanogenesis based on a genomic analysis[17]. The other three are affiliated to the hydrogenase group 4: two of the them (ANME-1a M5.MMPM and ANME-1 LC) with the group 4d, close to hydrogenases from *Thermoplasmata* and *Methanomicrobia* organisms, characterized as a multimeric complex that oxidize $H_2$ to reduce ferredoxin[24-27]; and the last hydrogenase (also from ANME-1 LC) with the group 4g, next to other euryarchaeotal hydrogenases that seem to couple ferredoxin oxidation to proton reduction[26].

Analysis of intraspecies distribution of hydrogenases within ANME-1c species

Given the apparent mosaic distribution of hydrogenases across ANME-1 lineages, we further detailed patterns of hydrogenase occurrence within the currently available genomes of ANME-1c. All MAGs of ´*Ca.* M. ujae´ and two out of five in ´*Ca.* M. jalkutatii´ (FW4382_bin126 and NA091.008_bin1) contained the hydrogenase operon, whereas the ´*Ca.* M. jalkutatii´ MAG FWG175, the most contiguous genome that was assembled into a single scaffold, does not contain hydrogenases. To confirm that the presence of hydrogenase genes in ´*Ca.* M. jalkutatii´ is different between MAGs, we mapped the metagenomic reads from our full South Pescadero Basin sample set to the MAGs. This analysis revealed that samples where ANME-1c MAGs did not have hydrogenase genes indeed did not have reads mapping the hydrogenase genes of MAGs FW4382_bin126 and NA091.008_bin1 (Fig. 3b). Additionally, the local absence of hydrogenase genes in FWG175 was confirmed in a genome-to-genome alignment (Fig. S4). Hydrogenase genes thus appear to be a part of the pangenomic repertoire of ´*Ca.* M. jalkutatii´. Since the presence of the hydrogenase operon varies even between subspecies (as demonstrated with ´*Ca.* M. jalkutatii´), hydrogenases might have been preserved in the ANME-1 pangenome as an environmental adaptation rather than as an absolute requirement for their methanotrophic core energy metabolism.

CRISPR-Cas system

The ANME-1 CRISPR repeats were frequently found to be directly associated with Type IB and Type III *cas* gene operons (see Fig. S6a for examples), typical in archaea. A 95% sequence identity cutoff indicates that the CRISPR repeats are shared by different ANME-1 lineages, yet different from the CRISPR repeats found in the '*Ca.* Alkanophagales', a sister group to the ANME-1 archaea. Surveying previously published and our newly assembled metagenomes from two hydrothermal vent systems in the Gulf of California, South Pescadero Basin[28,29] and this study (22 assemblies) and Guaymas Basin ([30], 13 assemblies, Supplementary Table 7), led to the extraction of 20649 unique ANME-1 CRISPR spacers. The CRISPR-mapped contigs were up to 80 kb in size and contained up to 532 unique protospacers (Fig.S6c). As shown in Fig. 4a, the ANME-1 MGEs from South Pescadero Basin were primarily targeted by spacers found locally (n=1912), but also showed a high number of matches to Guaymas Basin spacers (n=894).

Examples of Auxiliary Metabolic Genes (AMGs) encoded by ANME viruses.

Head-tailed and spindle-shaped viruses encode various proteins involved in nucleotide and amino acid metabolisms, including ribonucleoside triphosphate reductase (NrdD), Queuosine biosynthesis enzymes (QueCDEF), and asparagine synthase, which can respectively boost nucleotide, preQ, and amino acid synthesis (Fig. 5b, 5c). Tepeuvirus PBV144 encodes a phosphoenolpyruvate carboxykinase (PEPCK), a central component of the gluconeogenesis pathway; ahpuchvirus PBV299 encodes a H+/gluconate symporter (GntT). These enzymes may facilitate the carbon assimmilation of ANME-1 hosts (Supplementary Table 11). PhoU, involved in phosphate transport[31], and 3'-Phosphoadenosine-5'-phosphosulfate (PAPS) reductase, which activates sulfate for assimilation, were also detected in ekchuahviruses, suggesting a potential viral boost of cellular P and S intake in ANME-1 cellular hosts during infection. PAPS reductase has been previously found in bacterial and archaeal viruses in hypersaline and marine environments[32-34], while *phoU* was reported from pelagic metaviromic surveys in the Pacific Ocean, but not assigned to archaeal viruses[35]. The detection of these virus-encoded auxiliary metabolic genes (AMGs) from hydrothermal vent systems reflects a broader trend in phosphate and nutrient manipulation by viruses in diverse environmental settings.


1 White, R. H. Biosynthesis of coenzyme M (2-mercaptoethanesulfonic acid). *Biochemistry* **24**, 6487-6493 (1985).

2 Graham, D. E. & White, R. H. Elucidation of methanogenic coenzyme biosyntheses: from spectroscopy to genomics. *Natural Product Reports* **19**, 133-147, doi:10.1039/B103714P (2002).

3 White, R. H. Identification of an Enzyme Catalyzing the Conversion of Sulfoacetaldehyde to 2-Mercaptoethanesulfonic Acid in Methanogens. *Biochemistry* **58**, 1958-1962, doi:10.1021/acs.biochem.9b00176 (2019).

4 Graupner, M., Xu, H. & White, R. H. Identification of the Gene Encoding Sulfopyruvate Decarboxylase, an Enzyme Involved in Biosynthesis of Coenzyme M. *Journal of Bacteriology* **182**, 4862-4867, doi:doi:10.1128/JB.182.17.4862-4867.2000 (2000).

5 Albers, S.-V. & Jarrell, K. F. The archaellum: how Archaea swim. *Frontiers in Microbiology* **6**, doi:10.3389/fmicb.2015.00023 (2015).

6 Albers, S.-V. & Jarrell, K. F. The Archaellum: An Update on the Unique Archaeal Motility Structure. *Trends in Microbiology* **26**, 351-362, doi:https://doi.org/10.1016/j.tim.2018.01.004 (2018).

7 van Wolferen, M., Orell, A. & Albers, S.-V. Archaeal biofilm formation. *Nature Reviews Microbiology* **16**, 699-713 (2018).

8 Pohlschroder, M. & Esquivel, R. N. Archaeal type IV pili and their involvement in biofilm formation. *Frontiers in microbiology* **6**, 190 (2015).

9 Walker, D. *et al.* The archaellum of Methanospirillum hungatei is electrically conductive. . *mBio* **10**, e00579-00519 (2019).

10 Holmes, D. E., Zhou, J., Ueki, T., Woodard, T. & Lovley, D. R. Mechanisms for Electron Uptake by Methanosarcina acetivorans during Direct Interspecies Electron Transfer. *Mbio* **12**, e02344-02321 (2021).

11 Quemin, E. R. *et al.* First insights into the entry process of hyperthermophilic archaeal viruses. *J Virol* **87**, 13379-13385, doi:10.1128/jvi.02742-13 (2013).

12 Baquero, D. P. *et al.* in *Advances in Virus Research* Vol. 108 (eds Margaret Kielian, Thomas C. Mettenleiter, & Marilyn J. Roossinck) 127-164 (Academic Press, 2020).

13 Briegel, A. *et al.* Structural conservation of chemotaxis machinery across Archaea and Bacteria. *Environmental Microbiology Reports* **7**, 414-419, doi:https://doi.org/10.1111/1758-2229.12265 (2015).

14 Quax, T. E. F., Albers, S.-V. & Pfeiffer, F. Taxis in archaea. *Emerging Topics in Life Sciences* **2**, 535-546, doi:10.1042/ETLS20180089 (2018).

15 Li, Z., Rodriguez-Franco, M., Albers, S.-V. & Quax, T. E. F. The switch complex ArlCDE connects the chemotaxis system and the archaellum. *Molecular Microbiology* **114**, 468-479, doi:https://doi.org/10.1111/mmi.14527 (2020).

16 Meyerdierks, A. *et al.* Metagenome and mRNA expression analyses of anaerobic methanotrophic archaea of the ANME-1 group. *Environmental microbiology* **12**, 422-439, doi:doi:10.1111/j.1462-2920.2009.02083.x (2010).

17 Chadwick, G. L. *et al.* Comparative genomics reveals electron transfer and syntrophic mechanisms differentiating methanotrophic and methanogenic archaea. *PLOS Biology* **20**, e3001508, doi:10.1371/journal.pbio.3001508 (2022).

18 Zheng, K., Ngo, P. D., Owens, V. L., Yang, X.-p. & Mansoorabadi, S. O. The biosynthetic pathway of coenzyme F430 in methanogenic and methanotrophic archaea. *Science* **354**, 339-342, doi:10.1126/science.aag2947 (2016).

19 Michael, A. J. Polyamine function in archaea and bacteria. *Journal of Biological Chemistry* **293**, 18693-18701, doi:https://doi.org/10.1074/jbc.TM118.005670 (2018).

20 Morimoto, N. *et al.* Dual Biosynthesis Pathway for Longer-Chain Polyamines in the Hyperthermophilic Archaeon <i>Thermococcus kodakarensis</i>. *Journal of Bacteriology* **192**, 4991-5001, doi:doi:10.1128/JB.00279-10 (2010).

21 Kanehisa, M. & Goto, S. KEGG: Kyoto Encyclopedia of Genes and Genomes. *Nucleic Acids Research* **28**, 27-30, doi:10.1093/nar/28.1.27 (2000).

22 Mihara, H. & Esaki, N. Bacterial cysteine desulfurases: their function and mechanisms. *Applied Microbiology and Biotechnology* **60**, 12-23, doi:10.1007/s00253-002-1107-4 (2002).

23 Tchong, S.-I., Xu, H. & White, R. H. l-Cysteine Desulfidase: An [4Fe-4S] Enzyme Isolated from Methanocaldococcus jannaschii That Catalyzes the Breakdown of l-Cysteine into Pyruvate, Ammonia, and Sulfide. *Biochemistry* **44**, 1659-1670, doi:10.1021/bi0484769 (2005).

24 Porat, I. *et al.* Disruption of the Operon Encoding Ehb Hydrogenase Limits Anabolic CO<sub>2</sub> Assimilation in the Archaeon <i>Methanococcus maripaludis</i>. *Journal of Bacteriology* **188**, 1373-1380, doi:doi:10.1128/JB.188.4.1373-1380.2006 (2006).

25 Lie, T. J. *et al.* Essential anaplerotic role for the energy-converting hydrogenase Eha in hydrogenotrophic methanogenesis. *Proceedings of the National Academy of Sciences* **109**, 15473-15478, doi:10.1073/pnas.1208779109 (2012).
26 Greening, C. *et al.* Genomic and metagenomic surveys of hydrogenase distribution indicate H2 is a widely utilised energy source for microbial growth and survival. *The ISME Journal* **10**, 761-777, doi:10.1038/ismej.2015.153 (2016).
27 Søndergaard, D., Pedersen, C. N. S. & Greening, C. HydDB: A web tool for hydrogenase classification and analysis. *Scientific reports* **6**, 34212-34212, doi:10.1038/srep34212 (2016).
28 Wu, F. *et al.* Unique mobile elements and scalable gene flow at the prokaryote–eukaryote boundary revealed by circularized Asgard archaea genomes. *Nature Microbiology* **7**, 200-212, doi:10.1038/s41564-021-01039-y (2022).
29 Speth, D. R. *et al.* Microbial communities of Auka hydrothermal sediments shed light on vent biogeography and the evolutionary history of thermophily. *The ISME Journal*, doi:10.1038/s41396-022-01222-x (2022).
30 Dombrowski, N., Teske, A. P. & Baker, B. J. Expansive microbial metabolic versatility and biodiversity in dynamic Guaymas Basin hydrothermal sediments. *Nature Communications* **9**, 4999, doi:10.1038/s41467-018-07418-0 (2018).
31 Muda, M., Rao, N. N. & Torriani, A. Role of PhoU in phosphate transport and alkaline phosphatase regulation. *Journal of Bacteriology* **174**, 8057-8064, doi:10.1128/jb.174.24.8057-8064.1992 (1992).
32 Liu, Y. *et al.* Diversity, taxonomy, and evolution of archaeal viruses of the class Caudoviricetes. *PLOS Biology* **19**, e3001442, doi:10.1371/journal.pbio.3001442 (2021).
33 Mizuno, C. M. *et al.* Novel haloarchaeal viruses from Lake Retba infecting Haloferax and Halorubrum species. *Environmental Microbiology* **21**, 2129-2147, doi:https://doi.org/10.1111/1462-2920.14604 (2019).
34 Mara, P. *et al.* Viral elements and their potential influence on microbial processes along the permanently stratified Cariaco Basin redoxcline. *The ISME Journal* **14**, 3079-3092, doi:10.1038/s41396-020-00739-3 (2020).
35 Hurwitz, B. L., Brum, J. R. & Sullivan, M. B. Depth-stratified functional and taxonomic niche specialization in the 'core' and 'flexible' Pacific Ocean Virome. *The ISME Journal* **9**, 472-484, doi:10.1038/ismej.2014.143 (2015).
